# Slow oscillation-spindle coupling is negatively associated with emotional memory formation following stress

**DOI:** 10.1101/2020.06.08.140350

**Authors:** Dan Denis, Sara Y. Kim, Sarah M. Kark, Ryan T. Daley, Elizabeth A. Kensinger, Jessica D. Payne

## Abstract

Both stress and sleep enhance emotional memory. They also interact, with the largest effect of sleep on emotional memory being seen when stress occurs shortly before or after encoding. Slow wave sleep (SWS) is critical for long-term episodic memory, facilitated by the temporal coupling of slow oscillations and sleep spindles. Prior work in humans has shown these associations for neutral information in non-stressed participants. Whether coupling interacts with stress to facilitate emotional memory formation is unknown. Here, we addressed this question by reanalyzing an existing dataset of 64 individuals. Participants underwent a psychosocial stressor (32) or comparable control (32) prior to the encoding of 150-line drawings of neutral, positive, and negative images. All participants slept overnight with polysomnography, before being given a surprise memory test the following day. In the stress group, time spent in SWS was positively correlated with memory for images of all valences. Results were driven by those who showed a high cortisol response to the stressor, compared to low responders. The amount of slow oscillation-spindle coupling during SWS was negatively associated with neutral and emotional memory in the stress group only. The association with emotional memory was significantly stronger than for neutral memory within the stress group. These results suggest that stress around the time of initial memory formation impacts the relationship between slow wave sleep and memory.

## Introduction

Sleep aids in the consolidation of episodic memory (Stickgold, 2005; Payne, Ellenbogen, *et al.*, 2008; Diekelmann & Born, 2010; Rasch & Born, 2013). One of the main theoretical accounts of this process, the active systems consolidation theory, posits that across sleep, memories become less dependent on the hippocampus and more dependent on neocortical areas (Takashima *et al.*, 2006; Klinzing *et al.*, 2019). According to this theory, memory consolidation occurs primarily during periods of slow wave sleep (SWS) and is facilitated by the precise triple phase-locking of neocortical slow oscillations, thalamocortical sleep spindles, and hippocampal sharp-wave ripples (Rasch & Born, 2013; Klinzing *et al.*, 2019). Evidence for this triple-phase locking, and its importance for memory has emerged with rodents (Latchoumane *et al.*, 2017). In humans, this coordination has been demonstrated in epilepsy patients with intracranial hippocampal recordings (Staresina *et al.*, 2015). Although hippocampal ripples cannot be detected non-invasively, slow oscillation-spindle coupling as detected via scalp EEG has been associated with memory consolidation in healthy humans (e.g. Niknazar *et al.*, 2015; Mikutta *et al.*, 2019; Denis, Mylonas, *et al.*, 2020; Zhang *et al.*, 2020).

Notably, there is substantial evidence that some memories are more likely than others to be consolidated during sleep (Diekelmann *et al.*, 2009; Stickgold & Walker, 2013). Emotional valence acts as a prioritization cue in the selective consolidation of both negative (Payne, Stickgold, *et al.*, 2008; Nishida *et al.*, 2009; Payne *et al.*, 2015) and positive (Chambers & Payne, 2014a, 2014b; Kim et al., 2020) episodic memories (Payne & Kensinger, 2010; Lipinska *et al.*, 2019). Although much research has linked the consolidation of emotional memories to rapid eye movement (REM) sleep (Groch et al., 2013; Kim et al., 2019; Nishida et al., 2009; Payne et al., 2012; Payne, Stickgold, et al., 2008; Sopp et al., 2017; Wagner et al., 2001), newer work suggests SWS plays a role as well (Alger et al., 2018; Kim & Payne, 2020; Payne et al., 2015). In fact, several studies have found relationships between emotional memory and SWS but not REM sleep, raising the question of whether SWS and REM sleep differentially contribute to emotional memory consolidation (Benedict et al., 2009; Payne, 2011,, 2014; Payne et al., 2015; Wagner et al., 2002). The positive effects of SWS have often been shown in daytime naps rather than overnight designs (e.g. Alger et al, 2018; Payne et al, 2015), suggesting that sleep stages act differently on emotional memories depending on time of day (Alger *et al.*, 2018). Others suggest a complementary role for the two stages in emotional memory consolidation, whereby the SWS-REM cycles that naturally occur in overnight sleep serve to strengthen and integrate newly acquired emotional memory traces in preexisting memory networks (Cairney *et al.*, 2015).

In spite of these findings implicating SWS in emotional memory consolidation, the role of SWS-based oscillatory activity, including slow oscillation-spindle coupling, remains underexplored in regard to emotional memories. Research on slow oscillation-spindle coupling has focused almost exclusively on episodic memories for neutral information (though Latchoumane et al. (2015) employed a fear conditioning paradigm in rodents). A key distinction between neutral and emotional memory formation is the involvement of the amygdala (Payne & Kensinger, 2018). While theories of memory consolidation highlight dialogue between the hippocampus and neocortex during SWS as being involved in memory consolidation, interactions between the hippocampus and amygdala, and their relevance for behavior, is currently underexplored (though see Cox et al, 2020). Furthermore, many studies examining SWS in relation to memory focus exclusively on sleep stage correlations. Although broad sleep stage information provides insight into sleep’s role in memory consolidation, they fail to capture the neural activity that may more directly underlie consolidation processes during sleep. It is therefore important to consider both broad sleep stage macroarchitecture and the specific neurophysiological “micro” events when investigating the impact of sleep on memory.

Although recent theories (Hutchison & Rathore, 2015) and empirical studies (Kim et al., 2019; Nishida et al., 2009; Sopp et al., 2017) of REM sleep and emotional memory consolidation have examined EEG characteristics during REM sleep (e.g. REM theta (4-7Hz) activity), to our knowledge no studies in humans have assessed the role of slow oscillation-spindle coupling during SWS in emotional memory. Despite some research finding positive associations with sleep spindle activity and emotional memory consolidation (Kaestner *et al.*, 2013; Cairney *et al.*, 2014; Cellini *et al.*, 2016; Alger *et al.*, 2018) many other studies report no association (Prehn-Kristensen *et al.*, 2011; Baran *et al.*, 2012; Bennion *et al.*, 2015; Göder *et al.*, 2015; Payne *et al.*, 2015; Bolinger *et al.*, 2018). Given the likelihood that it is the *coupling* between spindles and slow oscillations that are important for consolidation processes, this may explain some of the mixed findings when assessing sleep spindles in isolation. Mechanistically, the coupling of spindles to slow oscillations has been suggested to be crucial to the induction of synaptic plasticity that underlies the formation of long-term memory representations in cortical networks (see Klinzing et al, 2019 for a recent review). Because research on the relationship between slow oscillation-spindle coupling and emotional memory does not yet exist, exploratory work assessing these associations is important if we are to more fully understand how emotional memories are consolidated during sleep. Exploring these relationships is one of our main goals here, as we attempt to better understand the relationship between features of SWS and emotional memory.

A second goal involves better understanding how stress interacts with sleep to benefit emotional memory (Bennion et al., 2015; Kim & Payne, 2020; Payne & Kensinger, 2018). When it comes to the selective processing of emotional memories, sleep is not the only state important for consolidation. Exposure to stress, and stress-related neuromodulators such as norepinephrine and cortisol, have been linked to better subsequent memory for emotional compared to neutral stimuli (Payne *et al.*, 2007; Shields *et al.*, 2017; Cunningham *et al.*, 2018). When these neuromodulators are present around the time of the initial encoding event, changes are triggered in brain regions relevant for emotional memory, including enhanced activity in and connectivity between the hippocampus, amygdala, and prefrontal cortex (PFC) (Veer *et al.*, 2011, 2012; Ghosh *et al.*, 2013; Vaisvaser *et al.*, 2013). Critically, these stress induced changes in neural activity and connectivity are associated with selective enhancement of emotional memory (Schwabe, 2017; Shields *et al.*, 2019).

Psychosocial stress has been shown to lead to alterations in subsequent sleep. In particular, one review highlighted a number of changes in sleep architecture following stress, including reductions in SWS and REM sleep duration (Kim & Dimsdale, 2007). More recently it has been shown that a post-stress nap shows reduced delta (0.5 - 4.5Hz) power compared to a control nap (Ackermann *et al.*, 2019). A small but growing body of work has shown sleep and stress interact to facilitate emotional memory formation (Payne & Kensinger, 2018; Kim & Payne, 2020 for review). Levels of cortisol during encoding have been positively correlated with subsequent memory after nocturnal sleep, but not after an equivalent period of daytime wake (Bennion *et al.*, 2015). This relationship is stronger for emotional memory compared to neutral memory (Bennion *et al.*, 2015). In a different analysis of the dataset reported on here, where stress prior to encoding was induced in half of the participants, theta oscillations (4-7Hz) during REM sleep predicted emotional memory in stressed participants, particularly those showing a high cortisol response following the stressor (Kim et al., 2019). These results suggest that elevations in stress-related neuromodulators during learning aid in the tagging of emotional memories, potentially via enhancement of amygdala-hippocampus-PFC connectivity, for preferential processing during sleep (Kim & Payne, 2020).

An important next step for understanding stress-sleep interactions in emotional memory is examining the potential impact of stress on oscillatory activity during SWS. Again, by investigating slow oscillation-spindle coupling, we can further understand how sleep physiology subserves not only memory for different types of encoding materials (neutral vs emotional memories), but also different types of encoding conditions (stressed or not). Here, we utilized a rich dataset containing a stress manipulation, overnight sleep with polysomnography, fMRI recordings (not utilized in the present report), and an emotional memory task. This dataset has previously been analyzed in studies of encoding-related changes in resting state functional connectivity in non-stressed control subjects (Kark & Kensinger, 2019a), effects of physiological arousal on emotional memory vividness (Kark & Kensinger, 2019b), repetition suppression effects in non-stressed control subjects (Kark *et al.*, 2020), and interactions between stress and theta activity during REM sleep (Kim et al., 2019). To answer our specific questions about interactions between stress and slow wave sleep, we conducted exploratory analyses on this dataset with the goal of answering the following questions:

1. Does time spent in SWS correlate with memory for emotional items (Payne *et al.*, 2015), neutral items (Groch *et al.*, 2015), or both emotional and neutral items?
2. Does slow oscillation-spindle coupling correlate with memory for neutral items, as in previous research (e.g. Mikutta et al., 2019), and does coupling also correlate with emotional memory?
3. Does stress exposure at the time of learning alter these relationships, perhaps by making it even more likely that emotional memories will be consolidated over neutral ones?

## Methods

### Participants

We analyzed data from a multi-day experiment designed to examine the effects of stress and sleep on emotional memory (see Kim et al., 2019; Kark & Kensinger, 2019 for other analyses of this dataset). In total, 65 participants (ages 18-31, M = 21.86, SD = 2.75) took part in the study. One participant was excluded due to a recording error. Participants were assigned to either the stress (n = 32, 19 female; M_age_ = 21.5 years, SD = 2.8) or control (n = 32, 15 female; M_age_ = 22.3 years, SD = 2.7) group. One participant from the control group was excluded from analysis of slow oscillation-spindle coupling due to an error with the PSG recording file. Participants reported no history of neurological, psychiatric, or sleep-related disorders, and were free of any other chronic medical conditions or medication affecting the central nervous system. Participants were compensated for their time. The study received ethical approval from the Boston College Institutional Review Board.

## Materials

### Trier Social Stress Task (TSST)

This task is a reliable, well-validated inducer of psychosocial stress. Participants in the stress group were given 10 minutes to prepare a five-minute speech on a topic (e.g. “Why are you the best candidate for a job?”) using information about themselves. They were told that they would give the speech to two judges and were given writing materials to prepare notes. At the end of the 10 minutes, participants were escorted to a separate room with two seated judges (confederates). Participants then had their notes taken from them and were asked to give the speech from memory. The confederate judges were instructed to keep neutral expressions throughout the speech. If participants completed the speech before the five minutes were up, they were told to continue. Following the speech, participants performed an arithmetic task aloud for five minutes (e.g. “Continuously subtract 13 from the number 1022 as quickly and accurately as possible”). If they made a mistake, they were told to restart from the beginning.

Participants in the control group performed a similar set of tasks, but designed to minimize psychosocial stress. They had 10 minutes to prepare for the speech task as well, but were informed they were in the control group to mitigate anticipatory stress. They were then escorted into the same room, but without recording equipment (though physiological measures were still taken). They then read their speech aloud in the empty room. After five minutes, they completed an arithmetic task in the empty room.

### Emotional memory task

Stimuli were 300 International Affective Picture System (IAPS) images and their corresponding line drawings (Kark & Kensinger, 2015). The use of line drawings was motivated by several reasons: 1) minimize confounds due to re-encoding during the recognition phase; 2) allow cueing of specific memories of the full color image without representing the images themselves; 3) minimize arousal effects during recognition by using retrieval cues that were less emotionally arousing than the images presented at encoding; 4) create a challenging recognition task with sufficient hit and false alarm rates for calculating memory scores.

Negative and positive images were preselected using the IAPS normative database for arousal and valence. Critically, negative and positive images were matched on arousal and absolute valence, and were both more arousing and more negative/positively valanced than neutral images (Kark & Kensinger, 2015).

### Encoding

During incidental encoding, participants viewed 150 negative (50), neutral (50), and positive (50) images. The images that were viewed, versus those held out as foils on the recognition task, were varied across participants. Each line drawing was presented for 1.5s, followed by the full color image for 3s. After viewing each image, participants indicated whether they would approach or back away from the scene if they encountered it in real life. This incidental encoding task facilitates deep encoding. Between each trial, a fixation cross was displayed for 6 - 12s (**Figure 1B**).

**Figure 1.**
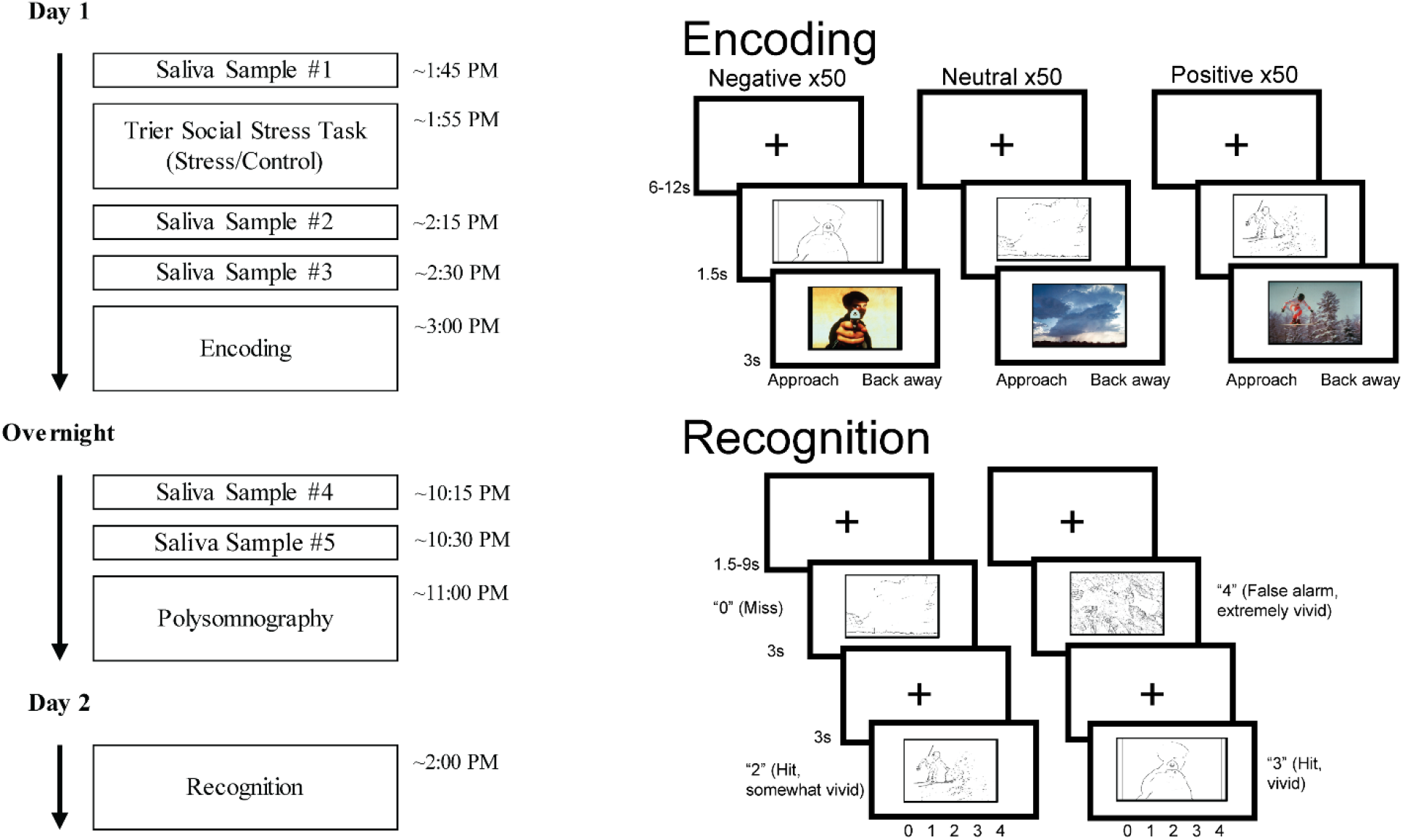
Study timeline and stimuli. **A**– Protocol details of relevance to the current analyses. On day 1, participants underwent the Trier Social Stress Task or control, then completed the encoding task. All participants then slept overnight in the lab with PSG recording. On day 2, participants completed a surprise recognition test. **B**-Encoding/retrieval task. During encoding (top), each line drawing was shown for 1.5s followed by the full IAPS image for 3s. Participants made an incidental encoding judgement (approach or back away) after each image pair. During recognition (bottom), each line drawing was shown for 3s, followed by an old-new judgement and vividness rating. IAPS = International Affective Picture System. Figure adapted from Kim et al. (2019). With permission from Wiley Periodicals, Inc.

### Recognition

Participants viewed only the line drawings during the recognition task. All 300 line drawings (100 negative, 100 neutral, 100 positive) were shown to every participant. One hundred fifty were the previously studied items, and 150 were new line drawings, with the encoding lists varying which items were to be classified into each of these categories. Each line drawing was presented for 3s. For each item, participants judged whether an item was new or old. If participants indicated that the item was old, they rated the vividness of their recollection on a scale of 0 = New, 1 = Old-not vivid, 2 = Old-somewhat vivid, 3 = Old-vivid, 4 = Old-Extremely vivid. Between trials, a fixation cross was displayed for 1.5 - 9s (**Figure 1B**).

### Procedure

For up to seven days prior to the experimental period, participants wore a wrist actigraphy monitor and kept a sleep log to monitor their sleep schedule. In the 24 hours before participation, participants were asked to refrain from caffeine, alcohol, and tobacco. To avoid contamination of cortisol samples, participants were told to refrain from physical activity, eating, drinking liquids aside from water, smoking, and brushing their teeth in the two hours prior to the study, and refrain from drinking water for at least 15 minutes prior to the study.

Upon arrival to the laboratory, participants provided a saliva sample for baseline cortisol levels (see Kim et al., 2019 for full details on saliva sampling or cortisol measurement). Participants were assigned to a group and completed either the stress (TSST) or control task. This was then followed by two further saliva samples taken approximately 15 minutes apart. (Additional saliva samples were taken later in the day to monitor circadian rhythms but will not be discussed further). Then, approximately 30 minutes after the TSST or control, they completed the encoding portion of the incidental emotional memory task during an fMRI scan. Participants then left the lab and went about their day before returning for overnight sleep monitoring that evening (approximately 6 hours after the end of the MRI session). Participants were instructed not to sleep in this intervening period. Two more saliva samples were taken prior to bedtime. All participants slept in the lab with polysomnographic recording. The following day (approximately 24 hours following encoding), participants completed a surprise recognition task, again in the MRI scanner (The study protocol as relevant to the current study is shown in **Figure 1A**. The full protocol is shown in **Supplementary Figure 1**).

### Polysomnography

Polysomnography (PSG) was acquired for a full night of sleep for all participants. PSG recordings included electrooculography recordings above the right eye and below the left eye, electromyography from two chin electrodes (referenced to each other), and electroencephalography (EEG) recordings from six scalp electrodes (F3, F4, C3, C4, O1, O2), referenced to the contralateral mastoids. Data were collected using a Grass Aura amplifier and TWin software at a sampling rate of 200Hz. Following acquisition, data were sleep scored in accordance with American Academy of Sleep Medicine (2007) guidelines. Data were artifact rejected using automated procedures in the Luna toolbox. Artifact free segments of data were subjected to further spectral analyses (see below). One participant had their sleep data removed due to a corrupted file.

## Sleep spectral analysis

### Sleep spindles

Spindles were detected at each electrode during SWS using a wavelet-based detector (Wamsley *et al.*, 2012). The raw EEG signal was convoluted with a 7-cycle complex Morlet wavelet with a peak frequency of 13.5Hz (full-width half-max bandwidth 12-15Hz) using the continuous wavelet transform implemented in MATLAB (*cwt* function). Spindle detection was performed on the squared wavelet coefficients after being smoothed with a 100ms moving average. A spindle was detected whenever the wavelet signal exceeded a threshold of nine times the median signal amplitude of artifact-free epochs for at least 400ms (Mylonas *et al.*, 2019). With these parameters, average spindle frequency was 13.18Hz.

### Slow oscillations

Slow oscillations were detected at each electrode during SWS using an automated algorithm. The data were band-pass filtered between 0.5 - 4Hz, and all positive-to-negative crossings were identified. Candidate slow oscillations were marked if two consecutive crossings fell 0.5 - 2 seconds apart. Peak-to-peak amplitudes for all candidate slow oscillations were determined, and oscillations in the higher 25th percentile (i.e. the 25% with the highest amplitudes) were retained and considered slow oscillations (Staresina *et al.*, 2015; Helfrich *et al.*, 2018).

### Slow oscillation-spindle coupling

Coupling was calculated at every electrode in SWS. The full, artifact free SWS signal was band-pass filtered in the delta (0.5 - 4Hz) and sigma (12 - 15Hz) bands. The Hilbert transform was applied to the whole artifact-free SWS signal to extract the instantaneous phase of the delta filtered signal and instantaneous amplitude of the sigma filtered signal. For each detected spindle, the peak amplitude of that spindle was determined. Then, it was determined whether the spindle peak occurred at anypoint during a detected slow oscillation (i.e., did the spindle peak fall between two consecutive positive-to-negative zero crossings which defined the start and end point of the slow oscillation). If a spindle was found to co-occur with a slow oscillation, the phase angle of the slow oscillation at the peak of the spindle was calculated. This approach to slow oscillation spindle coupling has been widely used elsewhere (e.g. Staresina *et al.*, 2015; Helfrich *et al.*, 2018; Mylonas *et al.*, 2020). We extracted the percentage of all spindles coupled with a slow oscillation at each electrode, the average phase of the slow oscillation at the peak of each coupled spindle, and the overall coupling strength (mean vector length). Coupling phase at each electrode was measured in degrees, with 0° indicating the spindle coupled at the positive peak of the slow oscillation. Coupling strength at each electrode was assessed using mean vector length measured on a scale of 0 −1, where 0 indicates each coupled spindle occurred at a different phase of the slow oscillation, and 1 indicates that each coupled spindle occurred at the exact same slow oscillation phase. For all statistical analyses, coupling values from frontal (F3, F4) and central (C3, C4) electrodes were averaged together.

### Statistical analysis

Behavioral results have been reported in detail elsewhere (Kim et al., 2019). Briefly, memory was measured as corrected recognition (hit rate - false alarm) for each valence type. Positive and negative stimuli were averaged together to create a combined emotional memory corrected recognition score. ANOVAs and follow up t-tests were used as appropriate. High cortisol responders and low responders were then identified via a median split (see Kim et al, 2019 for details), resulting in n = 16 high responders and n =16 low responders.

Relationships between sleep oscillatory measures and memory were assessed via multiple linear regression models. For all analyses, we looked both at positive + negative averaged together (hereafter referred to as emotional memory) as well as positive and negative separately. For all regression models, we were critically interested in the *interaction* between sleep measure (SWS time or coupling) and experimental group (stress or control). To control for multiple comparisons, interaction terms were evaluated at a false-discovery rate (FDR) adjusted significance level of *p* < .05. FDR adjustment was based on the four valence categories assessed (neutral, emotional (combined), positive, negative), and applied separately to analyses of SWS time and SWS coupling. Between-group comparisons of correlation coefficients were performed using Fischer’s r-to-z transformation (Fisher, 1925). Within-group comparisons were performed using Meng’s z test (Meng *et al.*, 1992). Both were carried out using the *cocor* package for R (Diedenhofen & Musch, 2015). For analyses of coupling, one participant from the stress group had their data excluded on the basis of being an outlier (>3SD from the mean). Appropriate circular statistics were used in all analyses involving coupling phase. Specifically, a Hotelling paired samples test was used to assess differences in coupling phase between frontal and central electrode sites (van den Brink *et al.*, 2014; van den Brink, 2020). Differences in coupling phase between the stress and control group were tested using a Watson-Williams test (Berens, 2009)., and correlations between coupling phase and memory were performed using circular-linear correlations (Berens, 2009).

## Results

### Behavior

Although behavioral results, including efficacy of the TSST in evoking stress, are reported in our previous publication (Kim et al., 2019), we summarize them briefly here (**Table 1; Figure 2**). When examining the effect of stress exposure on memory, we found there was a significant effect of valence (F (2, 124) = 4.69, *p* = .011, η_p_^2^ = .07). Memory at the recognition test was significantly better for both positive (*t* (63) = 2.46, *p* = .017, *d* = 0.31) and negative (*t* (63) = 2.77, *p* = .007, *d* = 0.35) images compared to neutral images. There was no difference between positive and negative items (t (63) = 0.38, *p* = .71, *d* = 0.05). A similar analysis was run for high and low cortisol responders within the stress condition (**Figure 2B**). There was a significant main effect of valence (F (2, 60) = 4.94, *p* = .010, η_p_^2^ = .014), with neutral items being more poorly remembered than both positive (t (31) = 2.34, *p* = .026, *d* = 0.41) and negative (t (31) = 2.83, *p* = .008, *d* = 0.50) items. For both analyses, there surprisingly was no effect of stress condition/reactivity, nor were there any significant interactions.

**Table 1.** Behavioral response data

|  | Hit rate |  | False alarm rate |  | d' |  |
| --- | --- | --- | --- | --- | --- | --- |
|  | Stress<br>M (SD) | Control<br>M (SD) | Stress<br>M (SD) | Control<br>M (SD) | Stress<br>M (SD) | Control<br>M (SD) |
| Neutral | 0.66 (0.11) | 0.67 (0.09) | 0.40 (0.17) | 0.41 (0.14) | 0.69 (0.46) | 0.66 (0.35) |
| Positive | 0.73 (0.11) | 0.74 (0.10) | 0.42 (0.18) | 0.46 (0.15) | 0.89 (0.59) | 0.73 (0.44) |
| Negative | 0.68 (0.10) | 0.69 (0.12) | 0.37 (0.18) | 0.42 (0.16) | 0.86 (0.50) | 0.68 (0.45) |

*Note.* M = mean, SD = standard deviation. Hits = proportion of trials correctly identified as old at the recognition test. False alarm = proportion of new trials incorrectly identified as old at the recognition test. $d' = d$ prime.

**Figure 2.**
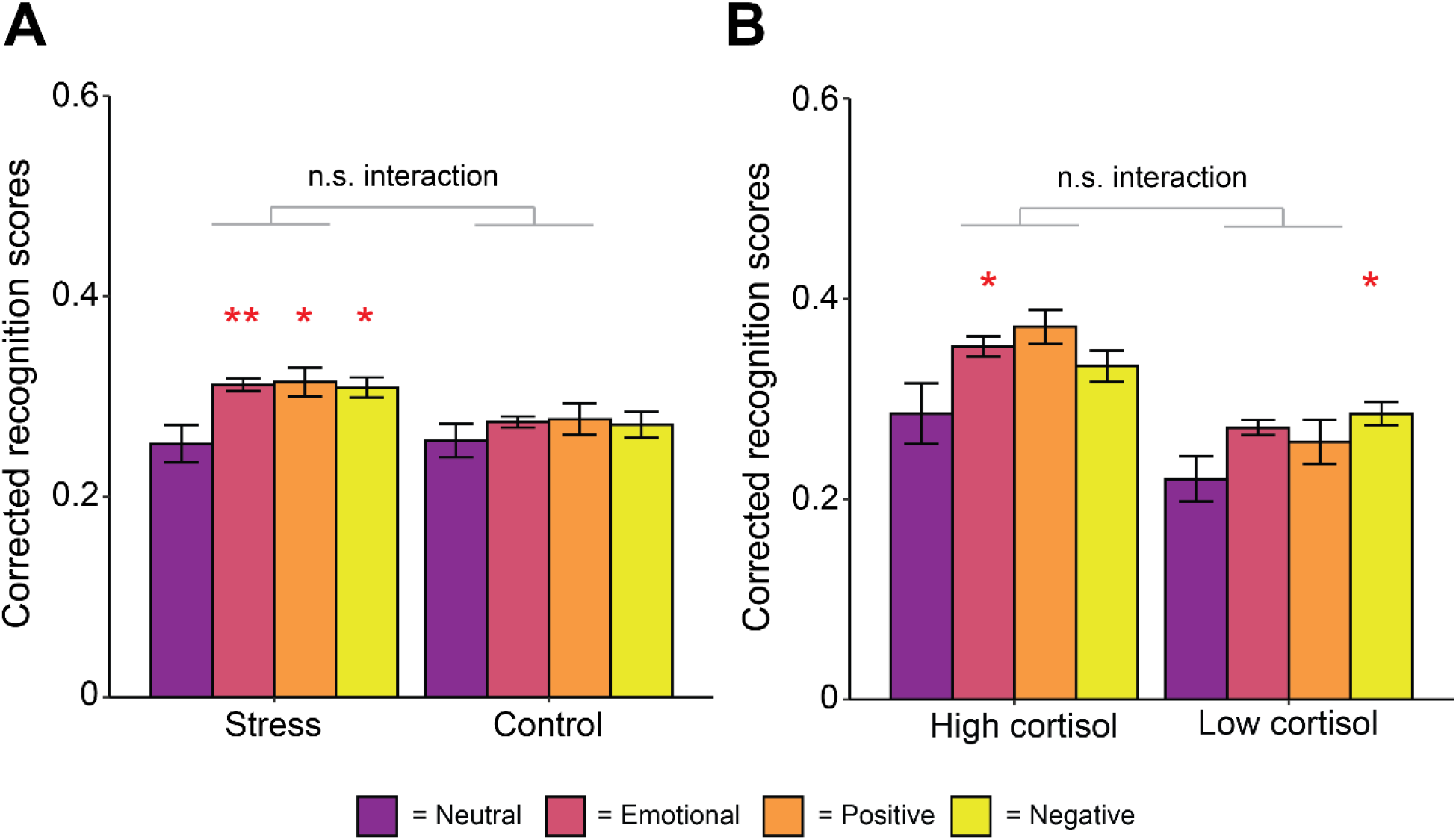
Behavior. Memory performance (hit rate – false alarm rate) for neutral, emotional (average of positive and negative), positive and negative items by **A**– experimental group and **B** – cortisol reactivity within the stress group, as determined by a median split. Errors bars represent the within-participant standard errors. Asterisks denote significance of exploratory within-group analyses of significant differences compared to neutral items ** = *p* < .01, * = *p* = .05.

In the stress group, the change in cortisol levels from pre- to- post TSST (maximum of the two post-TSST samples taken) significantly differed from zero (t (31) = 4.30, p < .001, d = 0.76). No such change was seen in the control group (t (31) = 0.94, p = .36, d = 0.17). The sample with the higher cortisol value was used as the peak post-TSST sample to better capture the slow time course of cortisol release and individual differences in this time course (de Kloet *et al.*, 2005). When we directly compared the stress and control groups on cortisol reactivity, reactivity was significantly higher in the stress group compared to the control group (t (38.97) = 3.72, p < .001, d = 0.93). However, elevated cortisol levels in the stress group did not persist into the evening. When participants returned to the lab to sleep, cortisol levels taken at bedtime (maximum of the two samples taken) were equivalent between the groups (t (42.85) = −1.58, p = .12, d = −0.40). Furthermore, levels of cortisol at bedtime were not significantly different to cortisol levels at the pre-stressor/control task sample (t (63) = −1.75, p = .08, d = −0.22).

## Sleep

### Group differences in sleep

Our previous report indicated no group differences in any measures of sleep macroarchitecture between the stress and control group (Kim et al, 2019). As a first step, we examined whether there were any group differences in SWS spectral measures. This would test whether the stressor (or stress reactivity) itself impacted SWS oscillatory activity during the sleep period. Results are displayed in **Table 2**. There were no significant differences between the stress and control group. Spindle and slow oscillation counts were significantly higher in low cortisol responders compared to high responders within the stress group, but these differences disappeared when stage time was controlled for using density measures (spindles/slow oscillations per minute of SWS).

**Table 2.** Group differences in SWS spectral measures

|  | Stress<br>M (SD) | Control<br>M (SD) | sig |
| --- | --- | --- | --- |
| Spindle count (n) | 303.5 (228) | 235.1 (181.2) | .19 |
| Spindle density (spindles/min) | 3.13 (1.78) | 2.55 (1.07) | .13 |
| Slow oscillation count (n) | 584 (243.3) | 596.1 (239.6) | .84 |
| Slow oscillation density (so/min) | 6.58 (0.91) | 6.97 (1.11) | .13 |
| Slow oscillation amplitude (uV) | 157.8 (26.25) | 164.8 (34.28) | .36 |
| % spindles coupled | 15.69 (6.52) | 16.14 (5.13) | .76 |
| Coupling strength (vector length) | 0.60 (0.15) | 0.60 (0.15) | .92 |
| Coupling phase angle (°) | 40.95 (19.01) | 35.61 (13.86) | .86 |
|  | High cortisol responders<br>M (SD) | Low cortisol responders<br>M (SD) | sig |
| Spindle count (n) | 209.3 (185.8) | 397.8 (232.3) | .017 |
| Spindle density (spindles/min) | 2.74 (1.56) | 3.53 (1.95) | .21 |
| Slow oscillation count (n) | 461.5 (232) | 706.6 (190.9) | .003 |
| Slow oscillation density (so/min) | 6.65 (1.18) | 6.51 (0.58) | .69 |
| Slow oscillation amplitude (uV) | 151.1 (27.88) | 164.5 (23.49) | .16 |
| % spindles coupled | 14.08 (4.81) | 17.31 (7.69) | .17 |
| Coupling strength (vector length) | 0.63 (0.13) | 0.58 (0.17) | .35 |
| Coupling phase angle (°) | 40.44 (11.17) | 41.48 (24.44) | .87 |

*Note.* M = mean, SD = standard deviation, sig = *p* value (uncorrected) from unpaired samples t-test. Differences in coupling phase assessed using a Watson-Williams two-sample test for circular data. so = slow oscillation. For coupling phase angle, 0° represents the peak of the spindle occurred at the positive peak of the slow oscillation.

### SWS time

There was a significant interaction between group (stress or control) and percentage of time spent in SWS for both neutral (β [95% confidence interval (CI)] = 0.51 [0.01, 1.00], *p* = .044) and emotional (combined) (β [95% CI] = 0.55 [0.07, 1.04], *p* = .026) items (see **Supplementary Table 1** for full regression model output). The same pattern was seen when positive (β [95% CI] = 0.48 [−0.01, 0.97], *p* = .054) and negative (β [95% CI] = 0.56 [0.07, 1.05], *p* = .025) items were examined separately. In the stress condition, a higher percentage of the sleep period spent in SWS was associated with better memory in all conditions (neutral, emotional, positive, and negative; all *r* > .35, all *p* < .043; **Figure 3**). However, this was not the case in the control group (all *r* > -.16, all *p* > .40). It should be noted that the overall regression models did not explain a significant proportion of the variance at the p < 0.05 level (**Supplementary Table 1**). We again underscore the preliminary and exploratory nature of our analyses, yet when adjusting for multiple comparisons using the false-discovery rate, all interaction terms approached, but did not meet, statistical significance (all *p_adj_* > .053).

**Figure 3.**
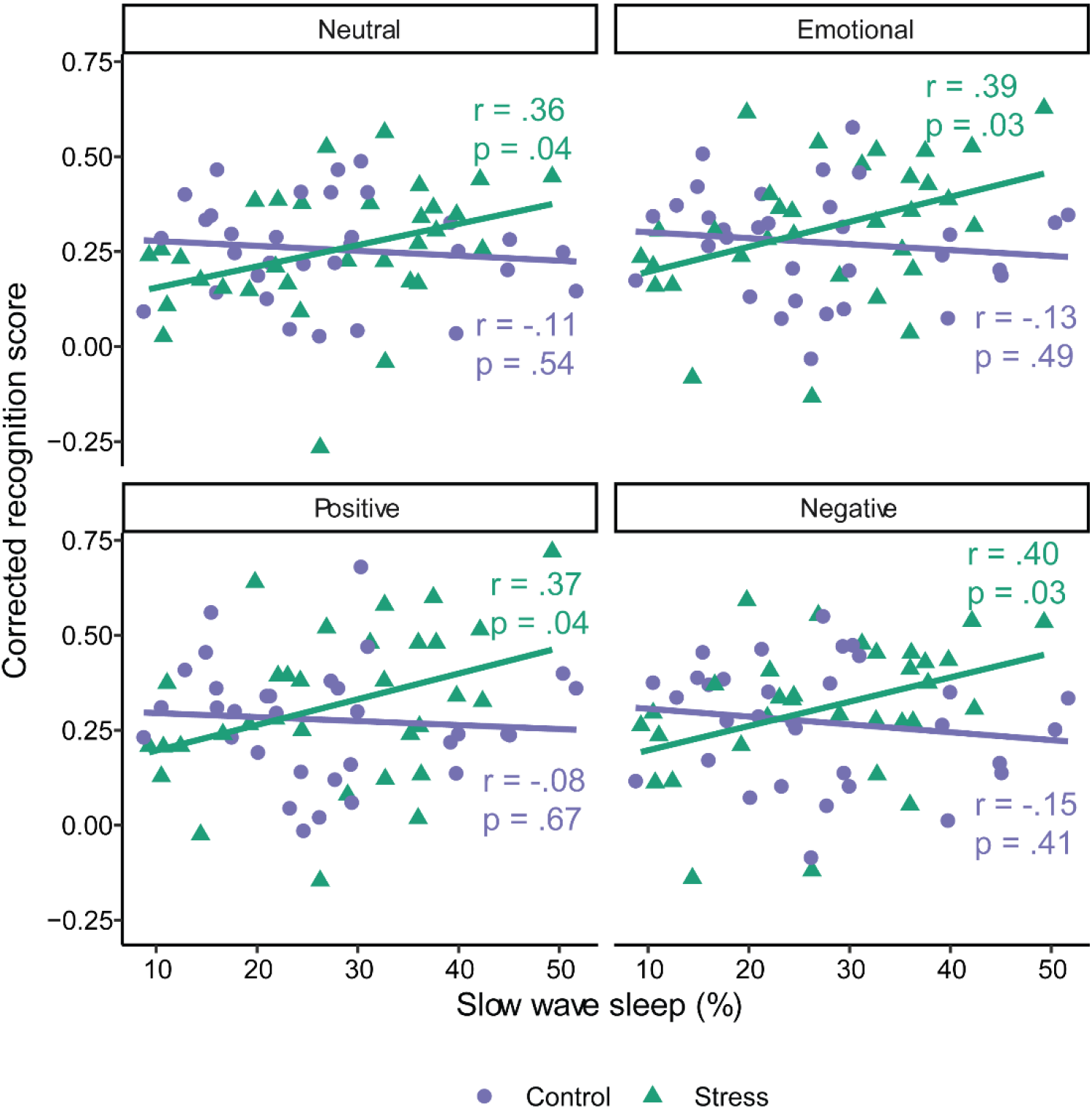
Associations between time spent in slow wave sleep and memory in the control and stress group. Emotional = Positive and negative item memory collapsed.

### SWS time in high and low cortisol responders

We next explored whether the relationship between SWS time and memory in the stress group was driven by those who showed a high cortisol response following the stressor. There was no significant interaction between cortisol response and SWS time for any valence type (neutral: β [95% CI] = −0.20 [−0.97, 0.57], p = 0.60; emotional (combined): β [95% CI] = −0.37 [−1.10, 0.35], p = 0.30; positive: β [95% CI] = −0.28 [−0.99, 0.43], p = 0.43; negative β [95% CI] = −0.45 [−1.20, 0.31], p = 0.24; **Figure 4; Supplementary Table 2**).

**Figure 4.**
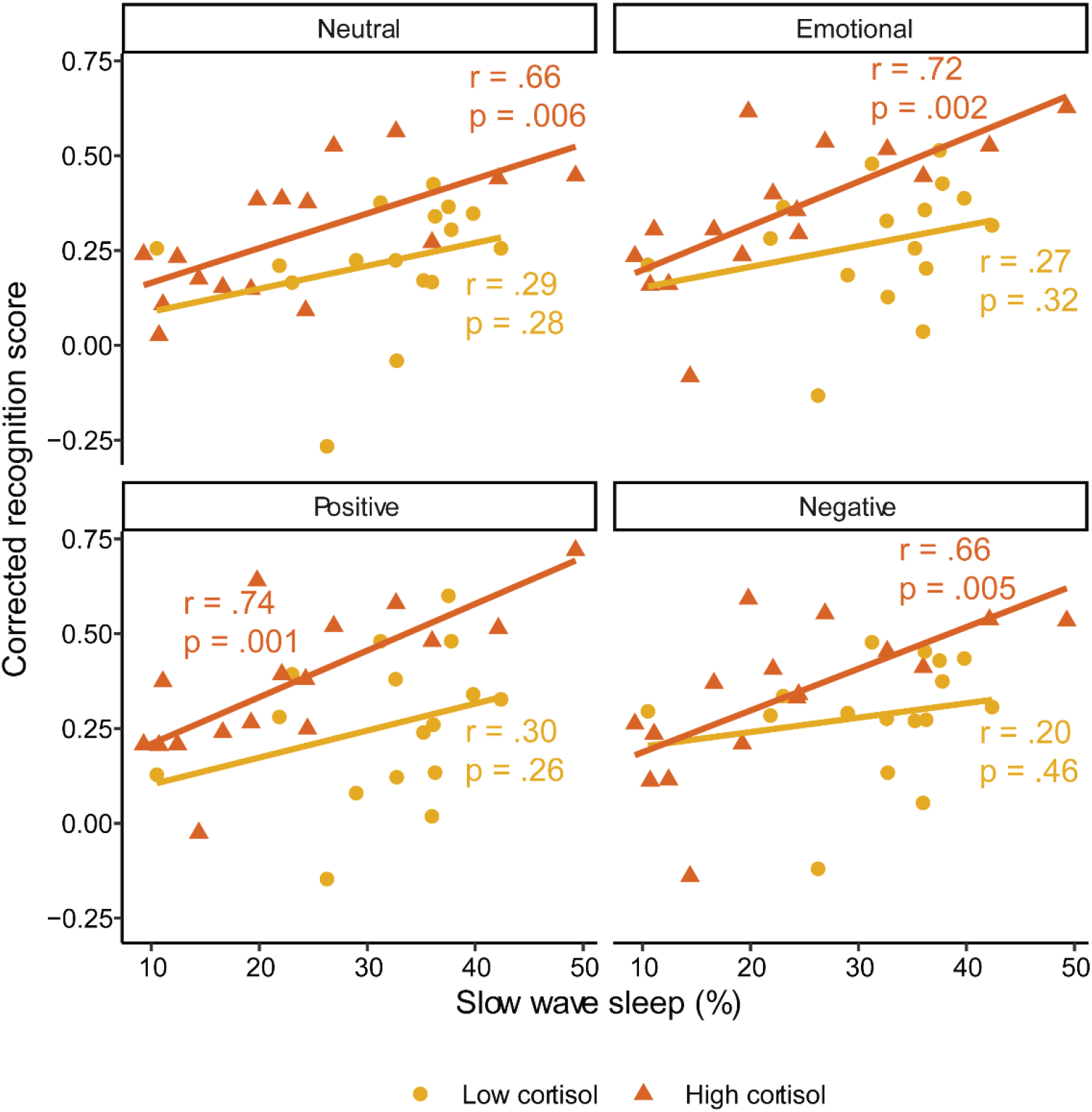
Associations between time spent in slow wave sleep and memory for low and high cortisol responders. Emotional = Positive and negative item memory collapsed.

Despite this, we note that correlations between SWS time and memory were only significant in the high responder group (neutral: *r* = .66, *p* = .006; emotional (combined): *r* = .72, *p* = .002; positive: *r* = .74, *p* = .001; negative: *r* = .66, *p* = .005). Correlations in the low responder group were all non-significant (all *r* < .31, *p* > .25). For all valence types, the between-group difference in correlation magnitude was not significant (all *z* < 1.63, *p* > .10).

### Slow oscillation-spindle coupling

Slow oscillation-spindle coupling dynamics during SWS are shown in **Figure 5**. When coupling phase was averaged across all coupled spindles within an individual, and across frontal and central electrodes, a Rayleigh test for non-uniformity was highly significant (*z* = 63, *p* < .001). This shows that across all participants, spindles showed a preferential phase coupling to the slow oscillation at M (SD) = 38.28° (16.88°) (**Figure 5A**). This corresponds to spindles preferentially coupling slightly after the peak of the slow oscillation (**Figure 5B**). Spindles coupled to slow oscillations later in the slow oscillation phase at frontal regions (M (SD) = 43° (17.55°)) compared to central regions (M (SD) = 32.95° (20.22°); F (2, 61) = 9.01, *p* < .001, **Figure 5**).

**Figure 5.**
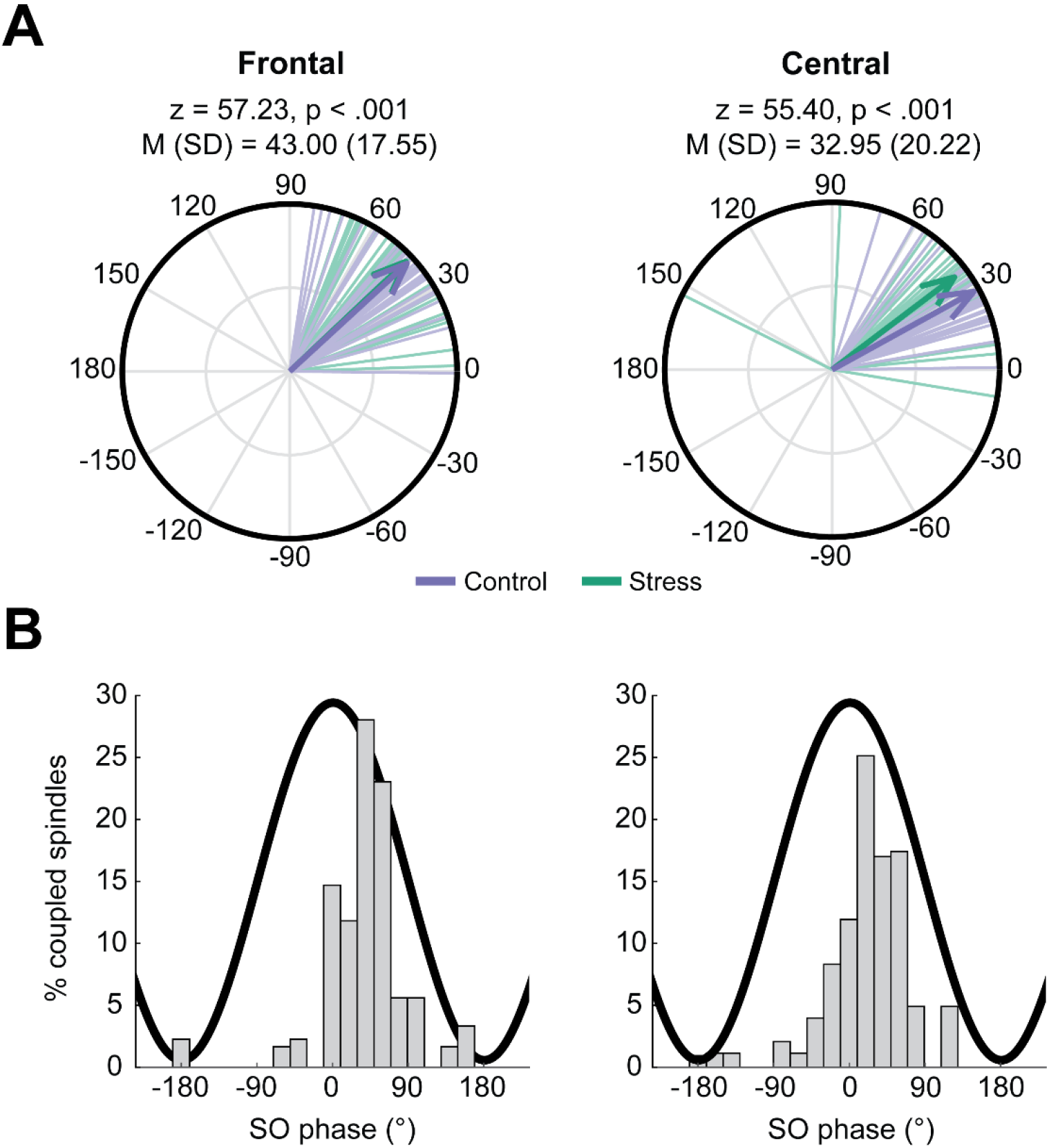
Slow oscillation spindle coupling during slow wave sleep. **A**– Circular phase plots displaying phase directions across participants at frontal (left plot) and central (right plot) electrode sites. Each line indicates the preferred coupling phase of individual participants. The direction of the arrow indicates the average phase across participants, with the length of the arrow indicating coupling strength across participants. Numbers above plot indicates result of Rayleigh tests for nonuniformity. **B –** Distribution of spindle coupling at different phases of the slow oscillation. We binned the percentage of spindles coupled to a slow oscillation (averaged across electrodes and participants) in 20° bins from −180° to 180°.

We investigated whether slow oscillation-spindle coupling during SWS was related to emotional memory following stress exposure. We focused specifically on coupling, due to emerging evidence that these events are specifically involved in memory processes (Klinzing *et al.*, 2019). First, we examined the overall amount of coupling during SWS. As such, the percentage of all SWS spindles that co-occurred with a slow oscillation was used as the dependent variable. We took the average of frontal (F3, F4) and central (C3, C4) electrodes. Full regression model results are shown in **Supplementary Table 3**.

There was a significant interaction between the amount of slow oscillation-spindle coupling and group (stress, control) for neutral (β [95% CI] = −0.69 [−1.18, −0.19], *p* = .007) and emotional (combined) (β [95% CI] = −0.69 [−1.14, −0.24], *p* =.003) memory. In both cases, this was driven by a significant *negative* relationship in the stress group (neutral: *r* = −.45, *p* = .01; emotional (combined): *r* = −.66, *p* < .001), meaning that higher amounts of coupling was associated with *worse* memory performance following stress (**Figure 6**). Importantly, the correlation with emotional memory was significantly larger than for neutral memory (*z* = 2.02, *p* = .04), suggesting that amount of coupling was more strongly related to impairment of emotional items compared to neutral. There were no significant relationships in the control group (neutral: *r* = .19, *p* = .30; emotional (combined): *r* = −.08, *p* = .69).

**Figure 6.**
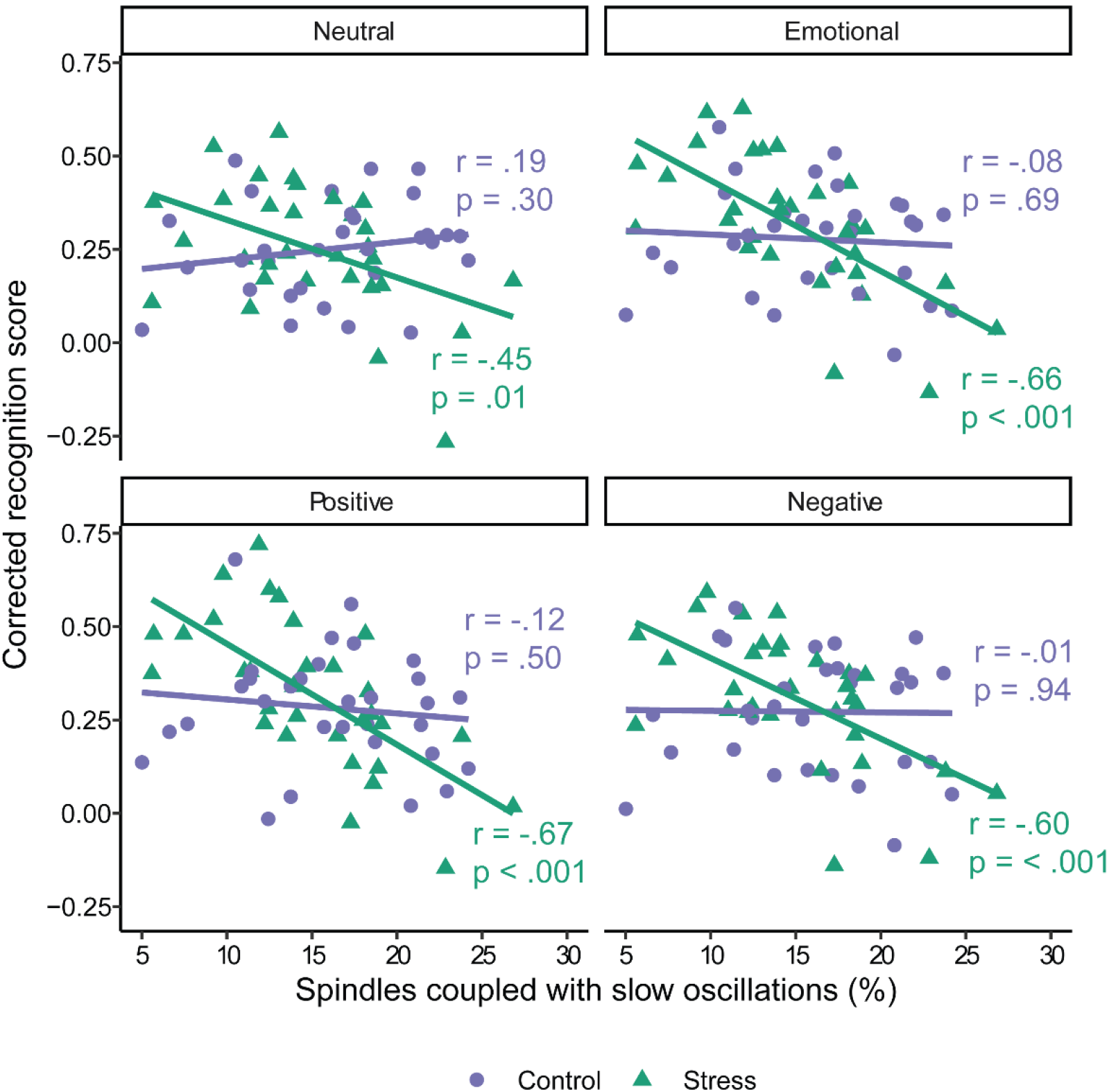
Associations between slow oscillation-spindle coupling and memory in the control and stress group. Coupling measures were averaged across frontal and central electrodes. Emotional = Positive and negative item memory collapsed.

Significant interactions between amount of coupling and stress condition were also found when positive (β [95% CI] = −0.66 [−1.10, −0.21], *p* = .004) and negative (β [95% CI] = −0.64 [−1.11, - 0.16], *p* = .01) items were examined separately. In both cases, a higher amount of coupling was associated with worse memory in the stress group (positive: *r* = −.67, *p* < .001; negative: *r* = −.60, *p* < .001), whereas there was no relationship in the control group (positive: *r* = −.12, *p* = .50; negative: *r* = −.01, *p* = .94). When adjusting for multiple comparisons using the false-discovery rate, all interaction terms remained significant (all *p_adj_* < .01).

### Coupling in high and low cortisol responders

Having found that the amount of slow oscillation-spindle coupling following stress is related to *worse* memory, we next sought to determine if the effect was driven by those who showed a high cortisol response to the stressor (**Figure 7**). Regression models did not indicate a significant interaction between amount of coupling and cortisol response for any valence type (neutral: β [95% CI] = −0.21 [−0.92, 0.50], p = 055; emotional (combined): β [95% CI] = −0.09 [−0.69, 0.50], p = .75; positive: β [95% CI] = −0.08 [−0.67, 0.50], p = .77; negative β [95% CI] = −0.10 [−0.74, 0.54], p = .75; **Supplementary Table 4**).

**Figure 7.**
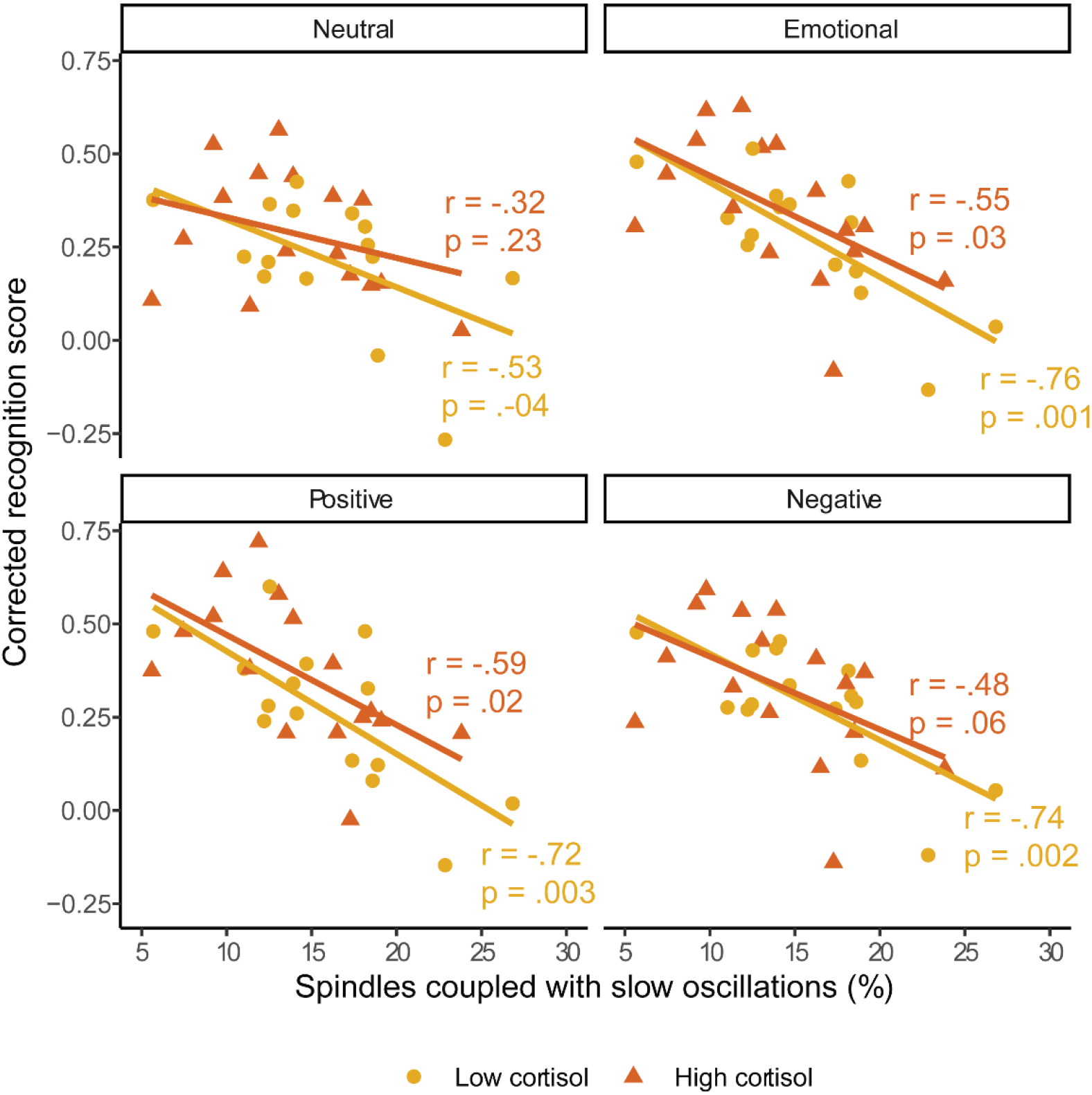
Associations between slow oscillation-spindle coupling in low and high cortisol responders. Coupling measures averaged across frontal and central electrodes. Emotional = Positive and negative item memory collapsed.

For emotional items, correlations between amount of coupling and emotional memory were significant or trending towards significant for both the high (emotional (combined): *r* = −.55, *p* = .03; positive: *r* = −.59, *p* = .02; negative: *r* = −.48; *p* = .06) and low responders (emotional (combined): *r* = −.76, *p* = .001; positive: *r* = −.72, *p* = .003; negative: *r* = −.74, *p* = .002). For neutral items, only low responders (*r* = −.53, *p* =.04) showed a significant correlation with memory (higher responders *r* = −.32, *p* = .23). For all valence types, the between-group difference in correlation magnitude was not significant (all *z* < 1.07, *p* > .29).

### Coupling phase and strength

Having shown that the amount of coupling is associated with impaired memory following stress, we next evaluated whether the timing and/or consistency of coupling events were also implicated in this effect. With regards to coupling strength, regression models for both neutral and emotional memory were non-significant (all R^2^ < .01, all *p* > .34). For coupling phase, circular linear correlations between coupling phase and memory were all non-significant (all *r* < .32, all *p* > .19). Together, these analyses suggest that the overall *amount* of slow oscillation-spindle coupling that is associated with worse memory following stress, and this effect is not related to the consistency or phase timing of the coupling events themselves.

### Sleep spindles and slow oscillations in isolation

Given the existing literature suggesting a critical role of slow oscillation-spindle coupling in memory processes (e.g. Klinzing *et al.*, 2019) we chose to primarily focus on coupling in this analysis. Nevertheless, we conducted supplemental analyses examining sleep spindle and slow oscillation densities in isolation. With regards to sleep spindles, all regression models were non-significant (all R^2^ < .04, all *p* > 18). With regards to slow oscillations, we did observe a pattern of results similar to coupling, in that a higher slow oscillation density was associated with worse memory in the stress group (see **supplemental results** for details). To ask whether coupling explained significantly more of the variance in emotional memory than slow oscillation density alone, we ran a multiple linear regression model predicting emotional memory from slow oscillation-spindle coupling and slow oscillation density. Coupling predicted emotional memory independently of slow oscillation density (β [95% CI] = −0.59 [−0.90, −0.27], *p* = .001). Slow oscillation density, however, did not predict emotional memory independently of coupling (β [95% CI] = −0.17, [−0.49, 0.15], *p* = .28). Adding coupling to the model significantly increased variance explained (adjusted R^2^ = .42) compared to slow oscillation density alone (adjusted R^2^ = .16), *F* (1, 28) = 14.28, *p* = .001.

### Additional post-hoc tests

There were no significant interactions with memory when coupling during stage 2 sleep was considered (all p > .44). Our primary metric of SWS coupling was the percentage of spindles coupled to slow oscillations. The reverse (percent of slow oscillations coupled to a spindle) did show a similar pattern of results, with a negative correlation between the amount of coupling and emotional memory in the stress (r = −.39) but not the control group (r = −0.039). When the number of coupled spindles (rather than the %) was considered, again a similar pattern of results were found, though results were generally weaker than when using the percentage. We note that the number of coupled spindles is confounded with the total number of spindles, with the two being extremely highly correlated (r = 0.90, p < .001). Comparatively, the percentage of spindles coupled was not significantly correlated with the total number of spindles (r = −0.16, p .20).

## Discussion

We set out to explore the role of slow wave sleep (SWS) and slow oscillation-spindle coupling during SWS in facilitating sleep-stress interactions on long-term emotional memory formation. Although recent theoretical and experimental evidence suggests a key role for SWS and slow oscillation-spindle coupling in memory consolidation (Klinzing *et al.*, 2019), their role in *emotional* memory consolidation has yet to be explored, and possible interactions with stress are unknown.

First, we queried whether time spent in SWS correlates with memory for emotional, neutral, or both emotional and neutral items. A large body of research has linked consolidation of neutral episodic memories with time spent in SWS (Plihal & Born, 1997; Ackermann & Rasch, 2014), and more recently this has been extended to the consolidation of emotional memory as well (Payne *et al.*, 2015; Alger *et al.*, 2018). Studies that directly compare the benefit of SWS on neutral and emotional memory have found mixed results, with some reporting correlations with just emotional memory (Payne *et al.*, 2015), or just neutral memory (Groch *et al.*, 2015). In the present study, we found that time spent in SWS was positively associated with memory for both neutral and emotional items, but only in stressed participants. This suggests a more general memory function for SWS time such that, following stress at least, more time spent in SWS provides a memory benefit, but this benefit occurs for all items equally, independent of their emotional valence. SWS time may facilitate a sleep-stress interaction on memory, but this is not specific for emotionally valanced material.

Next, we looked at slow oscillation-spindle coupling events, which are believed to mediate sleep-based memory processing (Rasch & Born, 2013; Klinzing *et al.*, 2019). Contrary to our SWS time finding, the amount of slow oscillation-spindle coupling was *negatively* correlated with memory in the stress group. This suggests that coupling impairs memory following stress, rather than promote it as is typically found for non-stressed participants and neutral memories (Niknazar *et al.*, 2015; Mikutta *et al.*, 2019). These effects were stronger for emotional items compared to neutral ones, suggesting that stress and SWS coupling may interact to impede emotional memory to a greater extent than neutral memory. This inverse relationship underscores the importance of considering both broad sleep stages and specific neurophysiological events when investigating sleep’s effect on memory. Following stress, it appears that sleep stage time and oscillatory events within that stage impact memory differently.

We did not find any associations between coupling phase and memory performance, contrary to other work which has suggested that fast spindles (~12-16Hz) coupled to the excitable slow oscillation upstate peak to be particularly important for memory (Helfrich *et al.*, 2018; Mikutta *et al.*, 2019; Muehlroth *et al.*, 2019). In our dataset, we found an average coupling phase of ~35°, suggesting preferential coupling just after the slow oscillation peak. Although this is similar to prior work using the same spindle detection parameters (Demanuele *et al.*, 2016), it is slightly later than other work in similar samples, who show preferential phase coupling just before or at the slow oscillation peak (Cox *et al.*, 2018; Helfrich *et al.*, 2018; Muehlroth *et al.*, 2019; Denis, Mylonas, *et al.*, 2020). Differences between sleep stages, differences in spindle definitions, and differences in algorithm may all in part explain these between study differences. Recent work has also suggested that fast spindles may be further subdivided into “early” fast (occur primarily on the rising phase of the slow oscillation) and “late” fast (occur primarily at or shortly after the slow oscillation peak) spindles (McConnell *et al.*, 2020). It is plausible that late fast spindles were primarily detected in this sample, especially given recent evidence that late fast spindles are more prevalent in SWS (McConnell *et al.*, 2020). Functional significance of these potential subtypes should be explored in future research.

Future research is needed to fully understand why slow oscillation-spindle coupling negatively impacts emotional memory following stress. One possibility is that the nature of memory consolidation during SWS is not immediately suitable for highly salient memories formed under stress. Slow oscillation-spindle coupling is believed to facilitate systems consolidation during SWS, whereby hippocampal-dependent memories become more dependent on neocortical sites and are integrated into existing knowledge networks (Rasch & Born, 2013; Klinzing *et al.*, 2019). This may not be appropriate for memories formed under stress where emotional tone is high. Over time, the affective response associated with a memory is diminished, leaving the experience itself with reduced emotional tone (Dolcos *et al.*, 2005). It has been suggested that REM sleep could be involved in this process (Walker & van der Helm, 2009), and it is possible that more REM-based emotional processing is needed before a memory becomes fully integrated in the cortex via SWS-based consolidation. Under this hypothesis, consolidation of emotional memories in the stress group during SWS may be damaging because they are undergoing systems consolidation before other necessary offline processing is achieved. This could be tested in future studies whereby both sleep and memory are measured over longer time periods. In sleep immediately following learning, coupling would be expected to be negatively associated with emotional memory as found here. However, over longer time periods this may be reversed as the memory becomes ready to be integrated more into cortical networks.

The role of the amygdala in emotional memory formation and its activity during REM sleep have been well documented (Murty *et al.*, 2010; Genzel *et al.*, 2015). Far less is known however about amygdala activity during SWS in humans. Recent research has shown that sharp-wave ripples occur in the human amygdala during SWS, and these are temporally linked with both ripples in the hippocampus and sleep spindles (though notably not slow oscillations) (Cox *et al.*, 2020). This provides a potential physiological basis for emotional memory consolidation during SWS, although an association with memory is yet to be reported. How amygdala-hippocampal interactions during SWS differ after stress is also unknown. As such, it remains possible that the negative association between coupling and emotional memory in the stress group could in part be driven by some currently uncharacterized process occurring between the hippocampus and the amygdala during SWS.

Why SWS time and SWS coupling show opposite associations with memory is difficult to reconcile and requires future work to fully understand. It is important to consider that coupling events are just one neurophysiological event that occurs during SWS and is a particularly rare event. Only ~15% of spindles couple to slow oscillations, and coupled spindle density has been reported as being around 0.5 per minute (Denis, Mylonas, *et al.*, 2020; Mylonas *et al.*, 2020). As such, coupling is a rare event in the broader SWS sleep state. It is possible then that while coupling events themselves impair memories formed under stress, other aspects of SWS may still be beneficial, and these are captured in a broad sense when considering just SWS time. For example, the neurochemical milieu of SWS differs greatly from wakefulness and promotes memory consolidation (Feld & Born, 2020).

We found that stress alters the association between SWS and memory. While the stressor significantly increased cortisol levels from baseline, and to a greater extent than the control task, these effects did not persist into the evening. When cortisol was measured before bedtime, there were no differences between the groups, and cortisol levels had returned to baseline levels. This suggests that the results were due to stress-related neuromodulatory activity at encoding, and not due to stressor-related cortisol results at bedtime.

It is notable that no associations between SWS activity and memory were found in the control group, given other research showing SWS-related benefits for both neutral and emotional episodic memory in non-stressed participants (Plihal & Born, 1997; Payne, Stickgold, *et al.*, 2008; Cox *et al.*, 2012; Payne, Chambers, *et al.*, 2012; Groch *et al.*, 2013; Ackermann & Rasch, 2014; Niknazar *et al.*, 2015; Payne *et al.*, 2015; Mikutta *et al.*, 2019; Denis, Mylonas, *et al.*, 2020). While the reason for this finding is unclear, one possible explanation may be the delay interval used in this study design. In non-stressed participants, several studies have shown the benefit of sleep is largest if learning occurs close before sleep, with a sleep effect not being detected when there is a long wake period between learning and sleep. This has been demonstrated for both neutral (Gais *et al.*, 2006; Talamini *et al.*, 2008; Payne, Tucker, *et al.*, 2012; Denis, Schapiro, *et al.*, 2020), and emotional memories (Payne, Chambers, *et al.*, 2012). In the present study, participants left the lab and went about their day before returning to sleep. During that interval, memory strength may have deteriorated to the point to which sleep was unable to act on them. On the other hand, in the stress group, heightened cortisol during encoding could have led to stronger initial representations that were tagged for further offline processing (Shields *et al.*, 2019), allowing later sleep to further strengthen and consolidate these memories.

Prior work has found mixed results with regards to sleep spindles and emotional memory. While some studies have found that SWS spindles do correlate with emotional memory consolidation (Cairney *et al.*, 2014; Alger *et al.*, 2018), other work has not found any association (Morgenthaler *et al.*, 2014; Sopp *et al.*, 2017; Bolinger *et al.*, 2018). Pharmacologically enhancing stage 2 spindles has also been documented to improve emotional memory (Kaestner *et al.*, 2013), although other correlational studies have shown no association between stage 2 spindles and emotional memory (Prehn-Kristensen *et al.*, 2011; Baran *et al.*, 2012; Göder *et al.*, 2015; Bolinger *et al.*, 2018). Similarly, we did not find stage 2 spindle coupling to relate to emotional memory in the present study. Clearly, more work is needed to untangle the exact role of sleep spindles on emotional memory. It is notable that no prior work (to our knowledge) has specifically examined slow oscillation-spindle coupling, which should be assessed in future research.

The exploratory nature of this work means that it is important for future studies to replicate these findings with *a priori* hypotheses. Additional limitations should also be considered. Although emotional memory recognition was numerically higher in the stress group compared to the control group, we did not detect a significant interaction effect between stress and memory. This may suggest the study was underpowered to detect this group level interaction, and future research should work to employ larger sample sizes. In addition, memory was only tested once. As such, it is not clear from this study whether sleep helped to facilitate a change in memory performance from pre- to post-sleep. This study also utilized a between-subjects design, making it more difficult and less well-powered to examine differences between stress and a non-stress control. The current design also utilized a different task to previous studies of sleep-stress interactions. Bennion and colleagues used the emotional memory trade-off task, which presents complex scenes and then uses a recognition memory test to separately assess memory for salient objects and the background scene on which they were presented (Bennion et al., 2015). This differs from the current task, which used line drawings to more holistically cue memory for complex scenes; the use of line drawings as cues is likely to lead to a harder retrieval task, and it also minimizes the presence of emotion in the retrieval cues; these design differences may have affected the pattern of results when compared to past designs (Bennion et al., 2015). Finally, this study examined the impact of sleep and stress on both positive and negative stimuli, as opposed to just negative stimuli as it Bennion et al., (2015). The present work also lacked a wake control group. As such, it is not possible to say that sleep led to a significant emotional memory benefit compared to time spent awake. This sleep-specific benefit, however, has been frequently documented in prior research (Payne, Stickgold, et al., 2008; Payne & Kensinger, 2010; though see Lipinska et al., 2019; Schäfer et al., 2020). Additionally, this does not preclude correlational analyses between memory and sleep neurophysiology, the primary focus of this investigation.

Notably, we did not find any differences in SWS oscillatory events between the stress and control groups. One notable exception was that the total number of spindles and slow oscillations were higher in low cortisol responders within the stress group than high cortisol responders. However, this difference disappeared when sleep stage time was controlled for by using density measure (spindles/slow oscillations per minute of SWS). This differs from previous research, which has found that psychosocial stress impacts the architecture of subsequent sleep, in particular reducing slow wave (0.5Hz-4.5Hz) oscillatory power (Ackermann *et al.*, 2019). There are several possible reasons for this discrepancy. First, previous work showing changes in slow wave power following stress were looking at changes within an afternoon nap that occurred in close proximity to the stressor, rather than overnight sleep (Ackermann *et al.*, 2019). Cortisol levels around the time of sleep onset and the sleep period itself may have been higher in the Ackermann et al. study than in ours which led to the alterations in spectral activity (Ackermann *et al.*, 2019). The natural circadian rhythm of cortisol means that cortisol levels around sleep onset in our overnight design (~11pm) would most likely be lower than at sleep onset in an afternoon nap. In addition, approximately 6 hours elapsed in our study between the stressor and sleep onset, as opposed to ~3 hours in the Ackermann et al study. Previous work has examined spectral power in a wide frequency range of 0.5-4.5Hz, which includes both slow oscillation and delta frequency activity. It is possible that stress leads to elevation in overall spectral power in this band without impacting the individually detected slow oscillatory events. Finally, Ackermann et al. noted that changes in sleep following stress were not seen across the whole nap period, but only in the first 15 minutes. How this time-dependent effect in a nap would map to a full night of sleep is unclear but is worthy of future investigation.

More generally, our findings demonstrate the importance of not relying solely on sleep stage correlations when probing sleep and memory associations (Lim *et al.*, 2020; Muehlroth & Werkle‐Bergner, 2020; Simor *et al.*, 2020). By simply assessing time spent in SWS, no consideration is given to specific neurophysiological events. It is equally important to also examine the specific oscillatory events that occur during different periods of sleep. Recent experimental work has emphasized a mechanistic role for discreet events occurring during SWS that underpin memory consolidation processes (Latchoumane *et al.*, 2017; Klinzing *et al.*, 2019). To this end, coupling between slow oscillations, sleep spindles, and sharp-wave ripples mediate the consolidation of memories during sleep (Latchoumane *et al.*, 2017). Furthermore, as we show here, these macro and micro level features of sleep can in fact have opposite effects on memory, and show selective effects based on memory valence and stress around the time of initial encoding.

In summary, this study adds to a small but growing body of research examining interactions between sleep and stress in the processing of emotional memories. We examined, for the first time, the role of key SWS oscillations in this process. To our surprise, we found that increased coupling between slow oscillations and sleep spindles impaired emotional memory following stress. Future research should seek to replicate this finding, and further probe the possibility that stress leads to a change in the function of slow oscillations and their coupling with spindles. If confirmed, this would have important implications for understand how sleep and stress facilitate long-term emotional memory.

## Data sharing statement

Data and analysis code needed reproduce the results reported in this manuscript are available via the Open Science Framework (Denis, 2021, https://osf.io/qeyrk/). Because the experimental stimuli came from a copyrighted source (International Affective Picture System - IAPS), it is not possible to freely share the stimuli. A list of IAPS images used is available alongside the data and analysis code.

## Author contributions

DD: Data analysis and writing of the manuscript, SYK: Writing of the manuscript and data analysis, SMK: Study design and data collection, RTD: Data collection, supervision of PSG acquisition, EAK: Funding acquisition, study design, supervision, JDP: Funding acquisition, study design, supervision. All authors contributed to the final version of the manuscript

## Acknowledgements

This work was supported by NSF Grant BCS 1539361 awarded to E.A.K and J.D.P, NIH shared instrumentation Grant S10OD020039 (Harvard Center for Brain Science, CBS), and NSF-GRFP DGE1258923 to S.M.K, pre-doctoral NRSE fellowship 5F31MH113304-02 to S.M.K, Sigmi Xi Grant-in-Aid of Research to S.M.K. We thank Leonore Bovy for assistance in data preparation and preliminary analyses, and Sara E. Alger for helpful comments on a draft of this manuscript. We thank Tala Berro, Kevin Frederiks, Sandry Garcia, Olivia Hampton, Lauren Lu, Yasmin Yacaby, and Stephanie Sherman for assistance with participant recruitment, data management, and TRIER data collection.

## Abbreviations

EEG: Electroencephalography
FDR: False discovery rate
fMRI: Functional magnetic resonance imaging
IAPS: International Affective Picture System
PSG: Polysomnography
PFC: Pre-frontal cortex
REM: Rapid eye movement
SWS: Slow wave sleep
TSST: Trier Social Stress Test

## References

Ackermann, S., Cordi, M., La Marca, R., Seifritz, E., & Rasch, B. (2019) Psychosocial Stress Before a Nap Increases Sleep Latency and Decreases Early Slow-Wave Activity. Front. Psychol., 10, 20.

Ackermann, S. & Rasch, B. (2014) Differential effects of non-REM and REM sleep on memory consolidation? Curr Neurol Neurosci Rep, 14, 430.

Alger, S.E., Kensinger, E.A., & Payne, J.D. (2018) Preferential consolidation of emotionally salient information during a nap is preserved in middle age. Neurobiology of Aging, 68, 34–47.

Baran, B., Pace-Schott, E.F., Ericson, C., & Spencer, R.M.C. (2012) Processing of emotional reactivity and emotional memory over sleep. J Neurosci, 32, 1035–1042.

Benedict, C., Scheller, J., Rose-John, S., Born, J., & Marshall, L. (2009) Enhancing influence of intranasal interleukin-6 on slow-wave activity and memory consolidation during sleep. FASEB J., 23, 3629–3636.

Bennion, K.A., Mickley Steinmetz, K.R., Kensinger, E.A., & Payne, J.D. (2015) Sleep and Cortisol Interact to Support Memory Consolidation. Cereb Cortex, 25, 646–657.

Berens, P. (2009) CircStat: A Matlab toolbox for circular statistics. Journal of statistical software, 31.

Bolinger, E., Born, J., & Zinke, K. (2018) Sleep divergently affects cognitive and automatic emotional response in children. Neuropsychologia, 117, 84–91.

Cairney, S.A., Durrant, S.J., Jackson, R., & Lewis, P.A. (2014) Sleep spindles provide indirect support to the consolidation of emotional encoding contexts. Neuropsychologia, 63, 285–292.

Cairney, S.A., Durrant, S.J., Power, R., & Lewis, P.A. (2015) Complementary roles of slow-wave sleep and rapid eye movement sleep in emotional memory consolidation. Cereb Cortex, 25, 1565–1575.

Cellini, N., Torre, J., Stegagno, L., & Sarlo, M. (2016) Sleep before and after learning promotes the consolidation of both neutral and emotional information regardless of REM presence. Neurobiol Learn Mem, 133, 136–144.

Chambers, A.M. & Payne, J.D. (2014a) The Influence of Sleep on the Consolidation of Positive Emotional Memories: Preliminary Evidence. AIMS Neuroscience, 1, 39–51.

Chambers, A.M. & Payne, J.D. (2014b) Laugh yourself to sleep: memory consolidation for humorous information. Exp Brain Res, 232, 1415–1427.

Cox, R., Hofman, W.F., & Talamini, L.M. (2012) Involvement of spindles in memory consolidation is slow wave sleep-specific. Learn. Mem., 19, 264–267.

Cox, R., Mylonas, D.S., Manoach, D.S., & Stickgold, R. (2018) Large-scale structure and individual fingerprints of locally coupled sleep oscillations. Sleep, 41, zsy175.

Cox, R., Rüber, T., Staresina, B.P., & Fell, J. (2020) Sharp-wave ripples in human amygdala and their coordination with hippocampus. bioRxiv, 2020.01.07.897413.

Cunningham, T.J., Leal, S.L., Yassa, M.A., & Payne, J.D. (2018) Post-encoding stress enhances mnemonic discrimination of negative stimuli. Learn. Mem., 25, 611–619.

de Kloet, E.R., Joëls, M., & Holsboer, F. (2005) Stress and the brain: from adaptation to disease. Nature Reviews Neuroscience, 6, 463–475.

Demanuele, C., Bartsch, U., Baran, B., Khan, S., Vangel, M.G., Cox, R., Hämäläinen, M., Jones, M.W., Stickgold, R., & Manoach, D.S. (2016) Coordination of Slow Waves with Sleep Spindles Predicts Sleep-Dependent Memory Consolidation in Schizophrenia. Sleep, 40, zsw013.

Denis, D. (2021) sleepLDF: SO-spindle coupling. Open Science Framework.

Denis, D., Mylonas, D., Poskanzer, C., Bursal, V., Payne, J.D., & Stickgold, R. (2020) Sleep spindles facilitate selective memory consolidation. bioRxiv, 2020.04.03.022434.

Denis, D., Schapiro, A.C., Poskanzer, C., Bursal, V., Charon, L., Morgan, A., & Stickgold, R. (2020) The roles of item exposure and visualization success in the consolidation of memories across wake and sleep. Learn. Mem., 27, 451–456.

Diedenhofen, B. & Musch, J. (2015) cocor: A Comprehensive Solution for the Statistical Comparison of Correlations. PLoS One, 10, e0121945.

Diekelmann, S. & Born, J. (2010) The memory function of sleep. Nat Rev Neurosci, 11, 114–126.

Diekelmann, S., Wilhelm, I., & Born, J. (2009) The whats and whens of sleep-dependent memory consolidation. Sleep Med Rev, 13, 309–321.

Dolcos, F., LaBar, K.S., & Cabeza, R. (2005) Remembering one year later: Role of the amygdala and the medial temporal lobe memory system in retrieving emotional memories. PNAS, 102, 2626–2631.

Feld, G.B. & Born, J. (2020) Neurochemical mechanisms for memory processing during sleep: basic findings in humans and neuropsychiatric implications. Neuropsychopharmacology, 45, 31–44.

Fisher, R.A. (1925) Statistical Methods for Research Workers. Oliver and Boyd, Edinburgh, Scotland.

Gais, S., Lucas, B., & Born, J. (2006) Sleep after learning aids memory recall. Learn Mem, 13, 259–262.

Genzel, L., Spoormaker, V.I., Konrad, B.N., & Dresler, M. (2015) The role of rapid eye movement sleep for amygdala-related memory processing. Neurobiology of Learning and Memory, REM Sleep and Memory, 122, 110–121.

Ghosh, S., Laxmi, T.R., & Chattarji, S. (2013) Functional Connectivity from the Amygdala to the Hippocampus Grows Stronger after Stress. J. Neurosci., 33, 7234–7244.

Göder, R., Graf, A., Ballhausen, F., Weinhold, S., Baier, P.C., Junghanns, K., & Prehn-Kristensen, A. (2015) Impairment of sleep-related memory consolidation in schizophrenia: relevance of sleep spindles? Sleep Med, 16, 564–569.

Groch, S., Wilhelm, I., Diekelmann, S., & Born, J. (2013) The role of REM sleep in the processing of emotional memories: evidence from behavior and event-related potentials. Neurobiol Learn Mem, 99, 1–9.

Groch, S., Zinke, K., Wilhelm, I., & Born, J. (2015) Dissociating the contributions of slow-wave sleep and rapid eye movement sleep to emotional item and source memory. Neurobiol Learn Mem, 122, 122–130.

Helfrich, R.F., Mander, B.A., Jagust, W.J., Knight, R.T., & Walker, M.P. (2018) Old Brains Come Uncoupled in Sleep: Slow Wave-Spindle Synchrony, Brain Atrophy, and Forgetting. Neuron, 97, 221–230.

Hutchison, I.C. & Rathore, S. (2015) The role of REM sleep theta activity in emotional memory. Front Psychol, 6, 1439.

Kaestner, E.J., Wixted, J.T., & Mednick, S.C. (2013) Pharmacologically Increasing Sleep Spindles Enhances Recognition for Negative and High-arousal Memories. Journal of Cognitive Neuroscience, 25, 1597–1610.

Kark, S.M. & Kensinger, E.A. (2015) Effect of emotional valence on retrieval-related recapitulation of encoding activity in the ventral visual stream. Neuropsychologia, 78, 221–230.

Kark, S.M. & Kensinger, E.A. (2019a) Post-encoding Amygdala-Visuosensory Coupling Is Associated with Negative Memory Bias in Healthy Young Adults. J. Neurosci., 39, 3130–3143.

Kark, S.M. & Kensinger, E.A. (2019b) Physiological arousal and visuocortical connectivity predict subsequent vividness of negative memories. NeuroReport, 30, 800–804.

Kark, S.M., Slotnick, S.D., & Kensinger, E.A. (2020) Forgotten but not gone: FMRI evidence of implicit memory for negative stimuli 24 hours after the initial study episode. Neuropsychologia, 136, 107277.

Kim, E.-J. & Dimsdale, J.E. (2007) The Effect of Psychosocial Stress on Sleep: A Review of Polysomnographic Evidence. Behavioral Sleep Medicine, 5, 256–278.

Kim, S.Y., Kark, S.M., Daley, R.T., Alger, S.E., Rebouças, D., Kensinger, E.A., & Payne, J.D. (2019) Interactive effects of stress reactivity and rapid eye movement sleep theta activity on emotional memory formation. Hippocampus, 1–13.

Kim, S.Y. & Payne, J.D. (2020) Neural correlates of sleep, stress, and selective memory consolidation. Current Opinion in Behavioral Sciences, 33, 57–64.

Klinzing, J.G., Niethard, N., & Born, J. (2019) Mechanisms of systems memory consolidation during sleep. Nat. Neurosci., 22, 1598–1610.

Latchoumane, C.-F.V., Ngo, H.-V.V., Born, J., & Shin, H.-S. (2017) Thalamic Spindles Promote Memory Formation during Sleep through Triple Phase-Locking of Cortical, Thalamic, and Hippocampal Rhythms. Neuron, 95, 424–435.

Lim, D.C., Mazzotti, D.R., Sutherland, K., Mindel, J.W., Kim, J., Cistulli, P.A., Magalang, U.J., Pack, A.I., de Chazal, P., & Penzel, T. (2020) Reinventing Polysomnography in the Age of Precision Medicine. Sleep Medicine Reviews, 52, 101313.

Lipinska, G., Stuart, B., Thomas, K.G.F., Baldwin, D.S., & Bolinger, E. (2019) Preferential Consolidation of Emotional Memory During Sleep: A Meta-Analysis. Front Psychol, 10, 1014.

McConnell, B.V., Kronberg, E., Teale, P.D., Fishback, G.M., Kaplan, R.I., Fought, A.J., Sillau, S.H., Dhanasekaran, A.R., Berman, B.D., Ramos, A.R., McClure, R.L., & Bettcher, B.M. (2020) The Aging Slow Wave: A Shifting Amalgam of Distinct Slow Wave and Spindle Coupling Subtypes Define Slow Wave Sleep Across the Human Lifespan. bioRxiv, 2020.05.28.122168.

Meng, X., Rosenthal, R., & Rubin, D.B. (1992) Comparing correlated correlation coefficients. Psychological Bulletin, 111, 172–175.

Mikutta, C., Feige, B., Maier, J.G., Hertenstein, E., Holz, J., Riemann, D., & Nissen, C. (2019) Phase-amplitude coupling of sleep slow oscillatory and spindle activity correlates with overnight memory consolidation. Journal of Sleep Research, 28, e12835.

Morgenthaler, J., Wiesner, C.D., Hinze, K., Abels, L.C., Prehn-Kristensen, A., & Göder, R. (2014) Selective REM-Sleep Deprivation Does Not Diminish Emotional Memory Consolidation in Young Healthy Subjects. PLOS ONE, 9, e89849.

Muehlroth, B.E., Sander, M.C., Fandakova, Y., Grandy, T.H., Rasch, B., Shing, Y.L., & Werkle-Bergner, M. (2019) Precise Slow Oscillation-Spindle Coupling Promotes Memory Consolidation in Younger and Older Adults. Sci Rep, 9, 1940.

Muehlroth, B.E. & Werkle‐Bergner, M. (2020) Understanding the interplay of sleep and aging: Methodological challenges. Psychophysiology, 57, e13523.

Murty, V.P., Ritchey, M., Adcock, R.A., & LaBar, K.S. (2010) fMRI studies of successful emotional memory encoding: A quantitative meta-analysis. Neuropsychologia, 48, 3459–3469.

Mylonas, D., Baran, B., Demanuele, C., Cox, R., Vuper, T.C., Seicol, B.J., Fowler, R.A., Correll, D., Parr, E., Callahan, C.E., Morgan, A., Henderson, D., Vangel, M., Stickgold, R., & Manoach, D.S. (2020) The effects of eszopiclone on sleep spindles and memory consolidation in schizophrenia: a randomized clinical trial. Neuropsychopharmacology, 1–9.

Mylonas, D., Tocci, C., Coon, W.G., Baran, B., Kohnke, E.J., Zhu, L., Vangel, M.G., Stickgold, R., & Manoach, D.S. (2019) Naps reliably estimate nocturnal sleep spindle density in health and schizophrenia. Journal of Sleep Research, 29, e12968.

Niknazar, M., Krishnan, G.P., Bazhenov, M., & Mednick, S.C. (2015) Coupling of Thalamocortical Sleep Oscillations Are Important for Memory Consolidation in Humans. PLOS ONE, 10, e0144720.

Nishida, M., Pearsall, J., Buckner, R.L., & Walker, M.P. (2009) REM sleep, prefrontal theta, and the consolidation of human emotional memory. Cereb Cortex, 19, 1158–1166.

Payne, J.D. (2011) Sleep on it!: stabilizing and transforming memories during sleep. Nature Neuroscience, 14, 272–274.

Payne, J.D. (2014) Seeing the Forest through the Trees. Sleep, 37, 1029–1030.

Payne, J.D., Chambers, A.M., & Kensinger, E.A. (2012) Sleep promotes lasting changes in selective memory for emotional scenes. Front Integr Neurosci, 6, 108.

Payne, J.D., Ellenbogen, J.M., Walker, M.P., & Stickgold, R. (2008) The role of sleep in memory consolidation. In Learning and Memory: A Comprehensive Reference: Vol. 2. Cognitive Psychology of Memory. Elsevier, Oxford, England, pp. 663–685.

Payne, J.D., Jackson, E.D., Hoscheidt, S., Ryan, L., Jacobs, W.J., & Nadel, L. (2007) Stress administered prior to encoding impairs neutral but enhances emotional long-term episodic memories. Learn. Mem., 14, 861–868.

Payne, J.D. & Kensinger, E.A. (2010) Sleep’s Role in the Consolidation of Emotional Episodic Memories. Curr Dir Psychol Sci, 19, 290–295.

Payne, J.D. & Kensinger, E.A. (2018) Stress, sleep, and the selective consolidation of emotional memories. Current Opinion in Behavioral Sciences, Emotion-cognition interactions, 19, 36–43.

Payne, J.D., Kensinger, E.A., Wamsley, E.J., Spreng, R.N., Alger, S.E., Gibler, K., Schacter, D.L., & Stickgold, R. (2015) Napping and the selective consolidation of negative aspects of scenes. Emotion, 15, 176–186.

Payne, J.D., Stickgold, R., Swanberg, K., & Kensinger, E.A. (2008) Sleep preferentially enhances memory for emotional components of scenes. Psychol Sci, 19, 781–788.

Payne, J.D., Tucker, M.A., Ellenbogen, J.M., Wamsley, E.J., Walker, M.P., Schacter, D.L., & Stickgold, R. (2012) Memory for semantically related and unrelated declarative information: the benefit of sleep, the cost of wake. PLoS ONE, 7, e33079.

Plihal, W. & Born, J. (1997) Effects of early and late nocturnal sleep on declarative and procedural memory. J Cogn Neurosci, 9, 534–547.

Prehn-Kristensen, A., Göder, R., Fischer, J., Wilhelm, I., Seeck-Hirschner, M., Aldenhoff, J., & Baving, L. (2011) Reduced sleep-associated consolidation of declarative memory in attention-deficit/hyperactivity disorder. Sleep Medicine, 12, 672–679.

Rasch, B. & Born, J. (2013) About sleep’s role in memory. Physiol Rev, 93, 681–766.

Schäfer, S.K., Wirth, B.E., Staginnus, M., Becker, N., Michael, T., & Sopp, M.R. (2020) Sleep’s impact on emotional recognition memory: A meta-analysis of whole-night, nap, and REM sleep effects. Sleep Medicine Reviews, 101280.

Schwabe, L. (2017) Memory under stress: from single systems to network changes. European Journal of Neuroscience, 45, 478–489.

Shields, G.S., McCullough, A.M., Ritchey, M., Ranganath, C., & Yonelinas, A.P. (2019) Stress and the medial temporal lobe at rest: Functional connectivity is associated with both memory and cortisol. Psychoneuroendocrinology, 106, 138–146.

Shields, G.S., Sazma, M.A., McCullough, A.M., & Yonelinas, A.P. (2017) The effects of acute stress on episodic memory: A meta-analysis and integrative review. Psychological Bulletin, 143, 636–675.

Simor, P., van der Wijk, G., Nobili, L., & Peigneux, P. (2020) The microstructure of REM sleep: why phasic and tonic? Sleep Medicine Reviews, 101305.

Sopp, M.R., Michael, T., Weeß, H.-G., & Mecklinger, A. (2017) Remembering specific features of emotional events across time: The role of REM sleep and prefrontal theta oscillations. Cogn Affect Behav Neurosci, 17, 1186–1209.

Staresina, B.P., Bergmann, T.O., Bonnefond, M., van der Meij, R., Jensen, O., Deuker, L., Elger, C.E., Axmacher, N., & Fell, J. (2015) Hierarchical nesting of slow oscillations, spindles and ripples in the human hippocampus during sleep. Nature Neuroscience, 18, 1679–1686.

Stickgold, R. (2005) Sleep-dependent memory consolidation. Nature, 437, 1272–1278.

Stickgold, R. & Walker, M.P. (2013) Sleep-dependent memory triage: evolving generalization through selective processing. Nat Neurosci, 16, 139–145.

Takashima, A., Petersson, K.M., Rutters, F., Tendolkar, I., Jensen, O., Zwarts, M.J., McNaughton, B.L., & Fernández, G. (2006) Declarative memory consolidation in humans: a prospective functional magnetic resonance imaging study. Proc. Natl. Acad. Sci. U.S.A., 103, 756–761.

Talamini, L.M., Nieuwenhuis, I.L.C., Takashima, A., & Jensen, O. (2008) Sleep directly following learning benefits consolidation of spatial associative memory. Learn Mem, 15, 233–237.

Vaisvaser, S., Lin, T., Admon, R., Podlipsky, I., Greenman, Y., Stern, N., Fruchter, E., Wald, I., Pine, D.S., Tarrasch, R., Bar-Haim, Y., & Hendler, T. (2013) Neural traces of stress: cortisol related sustained enhancement of amygdala-hippocampal functional connectivity. Front. Hum. Neurosci., 7, 313.

van den Brink, R.L. (2020) circt_htest.m. MATLAB Central File Exchange,.

van den Brink, R.L., Wynn, S.C., & Nieuwenhuis, S. (2014) Post-Error Slowing as a Consequence of Disturbed Low-Frequency Oscillatory Phase Entrainment. J. Neurosci., 34, 11096–11105.

Veer, I.M., Oei, N.Y.L., Spinhoven, P., van Buchem, M.A., Elzinga, B.M., & Rombouts, S.A.R.B. (2011) Beyond acute social stress: Increased functional connectivity between amygdala and cortical midline structures. NeuroImage, 57, 1534–1541.

Veer, I.M., Oei, N.Y.L., Spinhoven, P., van Buchem, M.A., Elzinga, B.M., & Rombouts, S.A.R.B. (2012) Endogenous cortisol is associated with functional connectivity between the amygdala and medial prefrontal cortex. Psychoneuroendocrinology, 37, 1039–1047.

Wagner, U., Fischer, S., & Born, J. (2002) Changes in emotional responses to aversive pictures across periods rich in slow-wave sleep versus rapid eye movement sleep. Psychosomatic Medicine, 64, 627–634.

Wagner, U., Gais, S., & Born, J. (2001) Emotional memory formation is enhanced across sleep intervals with high amounts of rapid eye movement sleep. Learn Mem, 8, 112–119.

Walker, M.P. & van der Helm, E. (2009) Overnight therapy? The role of sleep in emotional brain processing. Psychol Bull, 135, 731–748.

Wamsley, E.J., Tucker, M.A., Shinn, A.K., Ono, K.E., McKinley, S.K., Ely, A.V., Goff, D.C., Stickgold, R., & Manoach, D.S. (2012) Reduced sleep spindles and spindle coherence in schizophrenia: mechanisms of impaired memory consolidation? Biol Psychiatry, 71, 154–161.

Zhang, J., Yetton, B., Whitehurst, L.N., Naji, M., & Mednick, S.C. (2020) The Effect of Zolpidem on Memory Consolidation Over a Night of Sleep. Sleep, zsaa084.

