## Supplementary materials for "Slow oscillation-spindle coupling is negatively associated with emotional memory formation following stress" (from another version of this preprint: version 2, 2021-01-12)

Supplementary results

Slow oscillation density

There was a significant interaction between slow oscillation density and experimental group (stress or control) for neutral (β [95% CI] = -0.59 [-1.11, -0.07], *p* = .028) and emotional (combined) recognition memory (β [95% CI] = -0.78 [-1.27, -0.28], *p* = .003). For emotional items, this was driven by a significant negative correlation in the stress group (*r* = -.42, *p* = .016) but not the control group (*r* = .30, *p* = .10). The same pattern was true for neutral items, though individual correlation coefficients were not significant (stress: *r* = -.28, *p* = .12; control: *r* = .28, *p* = .13). When emotional items were broken down into positive and negative items, the same pattern of results was observed for both item types (positive: β [95% CI] = -0.70 [-1.21, -0.20], *p* = .007; negative: β [95% CI] = -0.76 [-1.26, -0.26], *p* = .004). All interactions remained significant after FDR adjustment (all p_adj_ < .029). The magnitude of the correlation between slow oscillation density and neutral and emotional items was not significantly different within the stress group (*z* = 1.20,  *p* = .23).

*Supplementary Table 1*. Full regression model results for SWS percentage analyses

| DV: Neutral items | *F* (3, 60) = 1.91, *p* = .14, R^2^ = .04 | |
| --- | --- | --- |
|  | β [95% CI] | *p* |
| Stress condition | -0.01 [-0.49, 0.48] | .984 |
| SWS percent | -0.09 [-0.43, 0.24] | .598 |
| Stress * SWS percent | 0.51 [0.01, 1.00] | .045 |
| DV: Emotional items | *F* (3, 60) = 2.59, *p* = .061, R^2^ = .07 | |
|  | β [95% CI] | *P* |
| Stress condition | 0.22 [-0.27, 0.70] | .371 |
| SWS percent | -0.11 [-0.44, 0.23] | .530 |
| Stress * SWS percent | 0.55 [0.07, 1.04] | .027 |
| DV: Positive items | *F* (3, 60) = 2.13, *p* = .11, R^2^ = .05 | |
|  | β [95% CI] | *P* |
| Stress condition | 0.20 [-0.29, 0.68] | .424 |
| SWS percent | -0.07 [-0.40, 0.27] | .703 |
| Stress * SWS percent | 0.48 [-0.01, 0.98] | .055 |
| DV: Negative items | *F* (3, 60) = 2.43, p = .07, R^2^ = .06 | |
|  | β [95% CI] | *p* |
| Stress condition | 0.21 [-0.27, 0.70] | .380 |
| SWS percent | -0.14 [-0.47, 0.20] | .420 |
| Stress * SWS percent | 0.56 [0.07, 1.05] | .025 |

Note. CI = Confidence intervals, R^2^= Adjusted R^2^, SWS = Slow wave sleep. For stress condition, the control group was the reference group.

*Supplementary Table 2*. Full regression model results for SWS percentage analyses in high and low cortisol responders

| DV: Neutral items | *F* (3, 28) = 3.57, *p* = .026, R^2^ = .20 | |
| --- | --- | --- |
|  | β [95% CI] | *p* |
| Cortisol response | -0.77 [-1.49, -0.06] | .036 |
| SWS percent | 0.59 [0.15, 1.02] | .011 |
| Cortisol * SWS percent | -0.20 [-0.97, 0.57] | .601 |
| DV: Emotional items | *F* (3, 28) = 5.21, *p* = .005, R^2^ = .29 | |
|  | β [95% CI] | *p* |
| Cortisol response | -0.85 [-1.53, -0.18] | .015 |
| SWS percent | 0.70 [0.29, 1.11] | .002 |
| Cortisol * SWS percent | -0.37 [-1.10, 0.35] | .304 |
| DV: Positive items | *F* (3, 28) = 5.65, *p* = .003, R^2^ = .31 | |
|  | β [95% CI] | *p* |
| Cortisol response | -0.99 [-1.65, -0.33] | .005 |
| SWS percent | 0.67 [0.26, 1.07] | .003 |
| Cortisol * SWS percent | -0.28 [-1.00, 0.43] | .428 |
| DV: Negative items | *F* (3, 28) = 3.94, *p* = .018, R^2^ = .22 | |
|  | β [95% CI] | *p* |
| Cortisol response | -0.63 [-1.34, 0.07] | .078 |
| SWS percent | 0.68 [0.25, 1.11] | .004 |
| Cortisol * SWS percent | -0.45 [-1.20, 0.31] | .240 |

Note. CI = Confidence intervals, R^2^ = Adjusted R^2^, SWS = Slow wave sleep. For cortisol response, low responders were the reference group.

*Supplementary Table 3*. Full regression model results for slow oscillation-spindle coupling analyses

| DV: Neutral items | *F* (3, 58) = 3.18, *p* = .030, R^2^ = .10 | |
| --- | --- | --- |
|  | β [95% CI] | *p* |
| Stress condition | -0.03 [-0.52, 0.46] | .893 |
| SO-SS coupling | 0.16 [-0.18, 0.51] | .346 |
| Stress * SO-SS coupling | -0.69 [-1.18, -0.19] | .008 |
| DV: Emotional items | *F* (3, 58) = 7.64, *p* < .001, R^2^ = .25 | |
|  | β [95% CI] | *p* |
| Stress condition | 0.14 [-0.31, 0.58] | .547 |
| SO-SS coupling | -0.06 [-0.38, 0.25] | .681 |
| Stress * SO-SS coupling | -0.69 [-1.14, -0.24] | .004 |
| DV: Positive items | *F* (3, 58) = 8.11, *p* < .001, R^2^ = .26 | |
|  | β [95% CI] | *p* |
| Stress condition | 0.11 [-0.33, 0.55] | .614 |
| SO-SS coupling | -0.11 [-0.42, 0.20] | .496 |
| Stress * SO-SS coupling | -0.66 [-1.10, -0.21] | .005 |
| DV: Negative items | *F* (3, 58) = 5.20, *p* = .003, R^2^ = .17 | |
|  | β [95% CI] | *p* |
| Stress condition | 0.14 [-0.33, 0.61] | .542 |
| SO-SS coupling | -0.01 [-0.34, 0.32] | .936 |
| Stress * SO-SS coupling | -0.64 [-1.11, -0.16] | .010 |

Note. CI = Confidence intervals, R^2^ = Adjusted R^2^, SO-SS= Slow oscillation-sleep spindle. For stress condition, the control group was the reference group

*Supplementary Table 4*. Full regression model results for slow oscillation-spindle coupling analyses in high and low cortisol responders

| DV: Neutral items | *F* (2, 27) = 2.65, *p* = .069, R^2^ = .14 | |
| --- | --- | --- |
|  | β [95% CI] | *p* |
| Cortisol response | -0.24 [-0.94, 9.45] | .478 |
| SO-SS coupling | -0.32 [-0.83, 0.19] | .210 |
| Cortisol * SO-SS coupling | -0.21 [-0.92, 0.50] | .545 |
| DV: Emotional items | *F* (2, 27) = 7.30, *p* < .001, R^2^ = .39 | |
|  | β [95% CI] | *p* |
| Cortisol response | -0.19 [-0.78, 0.39] | .504 |
| SO-SS coupling | -0.59 [-1.02, -0.17] | .009 |
| Cortisol * SO-SS coupling | -0.10 [-0.69, 0.52] | .747 |
| DV: Positive items | *F* (2, 27) = 8.06, *p* < .001, R^2^ = .41 | |
|  | β [95% CI] | *p* |
| Cortisol response | -0.30 [-0.88, 0.27] | .290 |
| SO-SS coupling | -0.60 [-1.20, -0.18] | .007 |
| Cortisol * SO-SS coupling | -0.08 [-0.67, 0.50] | .769 |
| DV: Negative items | *F* (2, 27) = 5.20, *p* = .006, R^2^ = .30 | |
|  | β [95% CI] | *p* |
| Cortisol response | -0.06 [-0.69, 0.57] | .852 |
| SO-SS coupling | -0.55 [-1.00, -0.09] | .022 |
| Cortisol * SO-SS coupling | -0.10 [-0.74, 0.54] | .752 |

Note. CI = Confidence intervals, R^2^ = Adjusted R^2^, SO-SS= Slow oscillation-sleep spindle. For cortisol response, low responders were the reference group.

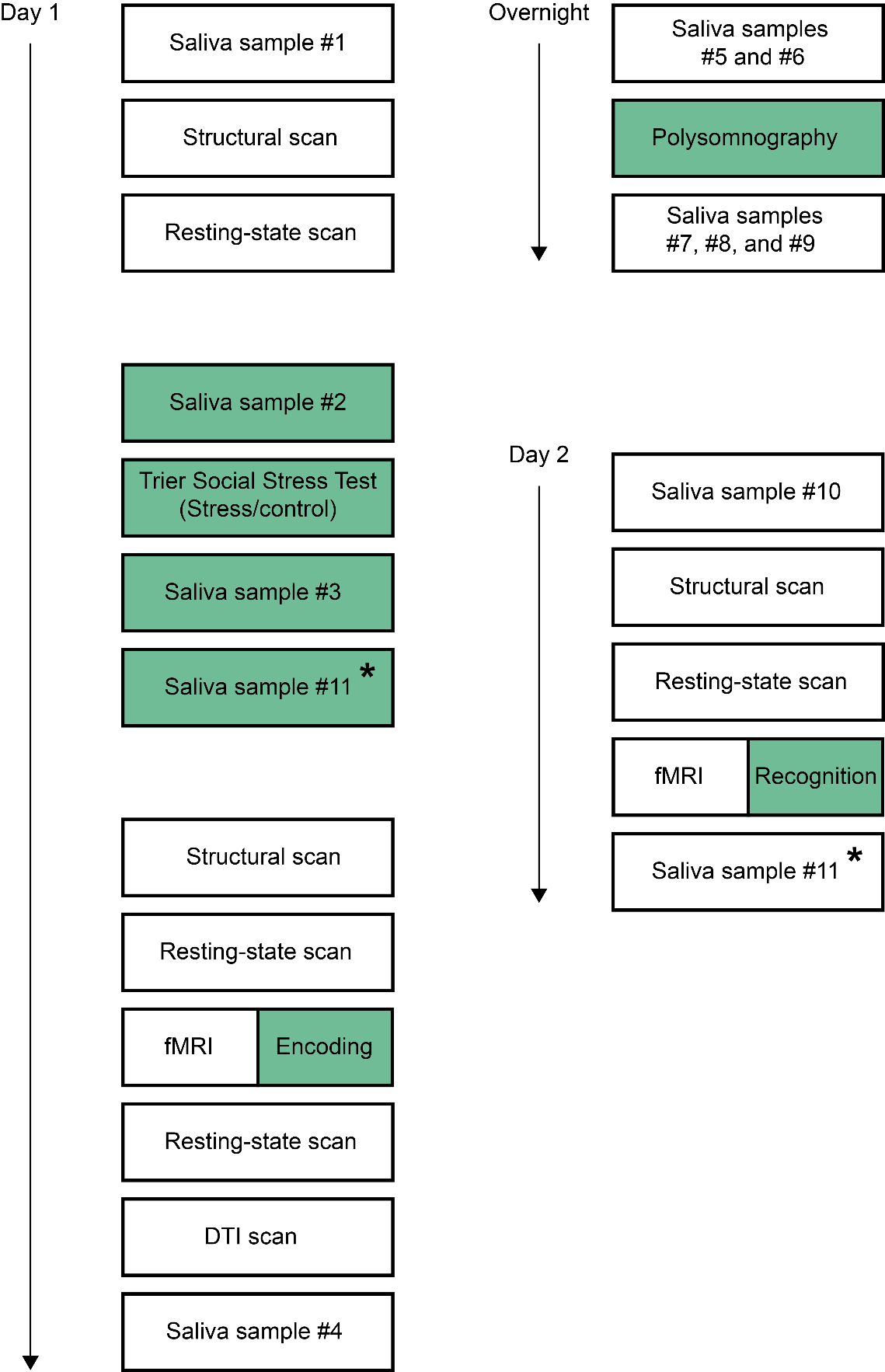

*Supplementary Figure 1*. Full experimental protocol. Note: Saliva sample #11 was taken after the recognition session on day 2 in the first n participants, after which the collection session moved to after the Stress/Control manipulation on day 1 (remaining n participants). Highlighted boxes indicate parts of the protocol reported on in this manuscript.
